# Structural reorganization of the chromatin remodeling enzyme Chd1 upon engagement with nucleosomes

**DOI:** 10.1101/089581

**Authors:** Ramasubramanian Sundaramoorthy, Amanda L. Hughes, Vijender Singh, Nicola Wiechens, Daniel P. Ryan, Hassane El-Mkami, Maxim Petoukhov, Dmitri I. Svergun, Barbara Treutlein, Monika Fischer, Jens Michaelis, Bettina Böttcher, David G. Norman, Tom Owen-Hughes

## Abstract

The yeast Chd1 protein acts to position nucleosomes across genomes. Here we model the structure of the Chd1 protein in solution and when bound to nucleosomes. In the apo state the DNA binding domain contacts the edge of the nucleosome while in the presence of the non-hydrolyzable ATP analog, ADP-beryllium fluoride, we observe additional interactions between the ATPase domain and the adjacent DNA gyre 1.5 helical turns from the dyad axis of symmetry. Binding in this conformation involves unravelling the outer turn of nucleosomal DNA and requires substantial reorientation of the DNA binding domain with respect to the ATPase domains. The orientation of the DNA-binding domain is mediated by sequences in the N-terminus and mutations to this part of the protein have positive and negative effects on Chd1 activity. These observations indicate that the unfavourable alignment of C-terminal DNA binding region in solution contributes to an auto-inhibited state.

## Introduction

Nucleosomes are not distributed randomly over the genomes of eukaryotes, but in general are aligned with respect to regulatory elements. In addition, the spacing between nucleosomes is also not random, with distinct nucleosome-to-nucleosome distances evident in different tissues and in different species (Hughes et al., 2012; van Holde, 1988). Abnormal inter-nucleosome spacing has been associated with increased intragenic transcription, and changes to nucleosome positioning at promoters results in changes to gene expression (Raveh-Sadka et al., 2012; Smolle et al., 2012). ATP-dependent chromatin remodelling enzymes act to organise nucleosomes over genomes with partial redundancy. For example, the Isw1 and Chd1 enzymes both play a major role in positioning nucleosomes over coding regions (Gkikopoulos et al., 2011; Ocampo et al., 2016; Radman-Livaja et al., 2012; Yen et al., 2012).

These enzymes belong to an extended family of Snf2-related chromatin proteins that can act to reconfigure DNA-protein interactions (Flaus et al., 2006), with many acting on nucleosomes. This family of proteins share an ATPase module comprised of two RecA-related domains capable of ATP-dependent DNA translocation (Havas et al., 2000; Lia et al., 2006; Saha et al., 2005; Zhang et al., 2006). The ATPase module is not found in isolation, but is associated with accessory domains both within the same polypeptide and as components of multi-subunit complexes. These accessory domains allow Snf2-related proteins to mediate different types of alteration to nucleosome structure or act on other protein-DNA complexes. For example, the Mot1 protein contains 16 HEAT repeats that facilitate engagement with the TBP protein (Wollmann et al., 2011).

Of all the Snf2 proteins, structural information is richest for the Chd1 proteins. The crystal structure of the ATPase domain in association with the N-terminal tandem chromodomains has been determined by crystallography (Hauk et al., 2010). In this structure, the chromodomains impede access to a putative DNA binding surface between the RecA domains, suggesting that reconfiguration is required in order for the RecA domains to engage productively with DNA. Supporting this, mutation of the chromodomains at the chromo-RecA interface increased the ATPase activity of Chd1 (Hauk et al., 2010). The concept that Snf2 proteins are subject to negative autoregulation is also supported by recent findings from the related ISWI proteins. Mutations in a region N-terminal to the ATPase domains of the *Drosophila* ISWI protein increase ATP-hydrolysis and remodelling activity (Clapier and Cairns, 2012). Interestingly the conformation of this region changes during ISWI action (Mueller-Planitz et al., 2013). The Chd1 protein has a C-terminal DNA binding domain (DBD) that is made up of SANT and SLIDE domains that are also present in ISWI proteins (Grune et al., 2003; Ryan et al., 2011). This second DNA binding interface is required for efficient nucleosome repositioning both in the context of Chd1 (Patel et al., 2012; Ryan et al., 2011; Sharma et al., 2011) and ISWI proteins (Dang and Bartholomew, 2007; Grune et al., 2003; Hota et al., 2013). Remarkably, substitution of this domain with a heterologous DNA – binding domain directs nucleosome positioning towards the DNA bound by this domain (McKnight et al., 2011; Patel et al., 2012).

Directed crosslinking has provided powerful insight as to the mode of interaction between remodelling enzymes and nucleosomes. Application of this approach to study the ISW2 enzyme showed that the ATPase domain engages with nucleosomal DNA near super helical location (SHL) 2, two turns from the dyad axis of symmetry. In the case of ISW2, the DNA binding accessory subunits are observed to engage linker DNA extending up to 50 bp from the edge of the nucleosome (Dang and Bartholomew, 2007; Kagalwala et al., 2004). In the case of ISWI containing enzymes it has been shown that two complexes can engage a single nucleosome (Racki et al., 2009) and this can facilitate the bidirectional movement of nucleosomes (Blosser et al., 2009). Single molecule fluorescence measurements have been used to monitor the transit of DNA through nucleosomes during the course of repositioning. These studies show that DNA is removed from nucleosomes in kinetically-coupled bursts of 3 bp that comprise of shorter single base increments (Deindl et al., 2013).

Existing structural information for chromatin remodelling enzymes is largely limited to subdomains. Less is known about the putatively unstructured regions connecting these domains and how these domains are oriented with respect to each other. Here we investigate the conformation of the ATPase Chd1 in solution and when engaged with nucleosomes. We find that there is a significant conformational change upon binding to nucleosomes. We obtain evidence to suggest that this change is limiting for Chd1 activity and contributes to maintenance of Chd1 in an auto-inhibited state. Regulation at this level provides a means of directing the action of remodelling ATPases towards specific aspects of nucleosome structure.

## Results

### Use of small angle x-ray scattering to study the solution structure of Chd1

We first sought to study the conformation of the Chd1 protein in solution. This is assisted by the fact that the structures of the chromoATPase domains and DNA binding domain (DBD) have been determined previously (Hauk et al., 2010; Ryan et al., 2011). The linkage between these domains however, is unclear as illustrated in Figure 1A. To help characterise the structure of intact Chd1 a series of fragments of Chd1 were expressed and purified (Figure 1B; Figure 1 – figure supplement 1). We then collected small angle X-ray scattering (SAXS) data for each of these (Figure 1 – figure supplement 2). For each fragment the hydrodynamic radius of the protein fragment and molecular weight in solution was calculated (Figure 1B). The values obtained are consistent with Chd1 being predominantly monomeric in solution. Volumes consistent with each scattering curve were calculated *ab initio* (Figure 1D). These volumes are consistent with the known structural features of Chd1 (Figure 1A). For example, the DBD can be docked within the volume obtained for this fragment of the protein and the chromoATPase domains within the volumes obtained for fragments that include this region. The volumes for the smaller fragments can be arranged within those of the larger fragments (Figure 1D). This indicates that the DBD and N-terminal 133 residues contribute to the protrusion adjacent to one of the ATPase domains (Figure 1D).

**Figure 1.**
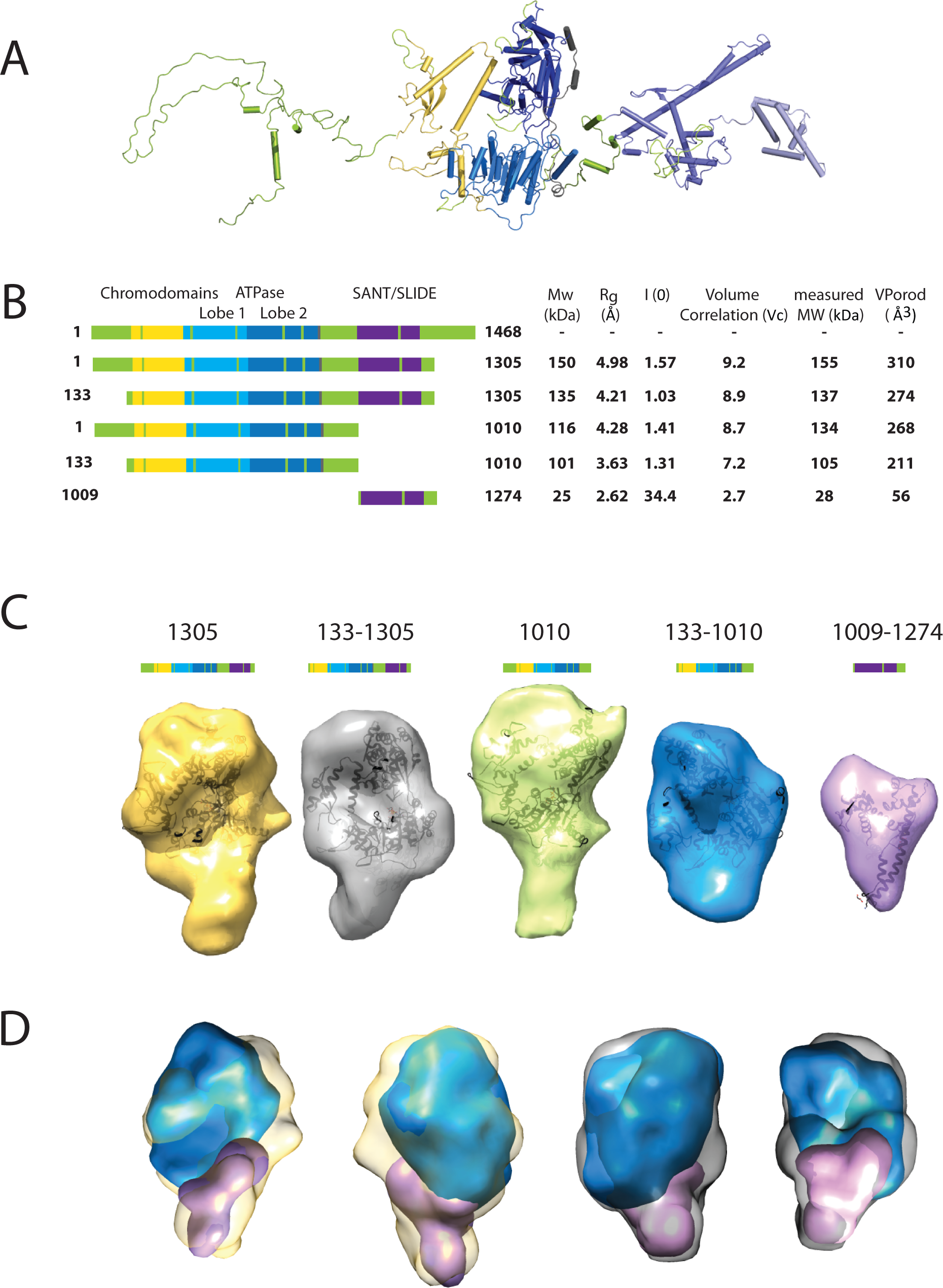
Characterising the solution structure of Chd1 by Small Angle X-ray scattering (SAXS). **A**, The overall structure of *S. cerevisiae* Chd1 molecule. Previously characterised structural features including the chromoATPase (3MWY) (Hauk et al., 2010) with chromodomains coloured yellow, ATPase lobe 1 - marine, ATPase lobe 2 – blue. The DNA binding domain (2XB0) (Ryan et al., 2011) is coloured deep blue with the NMR structure of the C-terminal extension (2N39) (Mohanty et al., 2016) in pale blue. The unresolved structural elements are coloured pale green and a cartoon representation of their predicted secondary structure provided to give an idea of scale. **B**, On the left Illustration of various Chd1 truncations. The known domains within the truncation are labelled and coloured. On the right the data (hydrodynamic radius (Rg), extrapolated zero intensity (I(0)), molecular weight (MW)) obtained from the SAXS analysis of the respective construct protein. **C**, Ab-initio bead models generated from the one-dimensional scattering curves using GASBOR for different construct. Known Chd1 crystal structures are docked into the respective volume. **D**, The volumes for 133-1010 (blue) and 1009-1274 (pink) are fitted into 1305 (yellow) and 133-1305 (grey) volume maps using SASREF.

### A structural model for Chd1 based on pulsed EPR measurements

The volumes obtained using SAXS are limited by the resolution possible using this approach. To obtain higher resolution models we used site directed spin labelling and pulsed electron-electron double resonance (PELDOR) measurements to characterise the Chd1 protein. To do this we first removed the six native cysteine residues converting them to serine. Importantly, this cys-free mutant protein displays nucleosome remodelling activity comparable to the wild type protein (Figure 2 – figure supplement 1). Pairs of cysteine residues were then introduced at specific sites in Chd1 via site-directed mutagenesis and these sites were then labelled with the thiol-reactive reporter (1-Oxyl-2,2,5,5-tetramethylpyrroline-3-methyl) methanethiosulfonate (MTSL). The interaction between attached MTSL groups can be measured by PELDOR and used to extract distance information (Pannier et al., 2000). To validate our approach, we first probed residues within the chromoATPase domains for which there is good structural information (Figure 2 – figure supplement 2). The raw dipolar evolution signal for each pair of labelled sites was subject to background correction and Tikhonov regularisation to obtain a distance distribution describing the positions of the labels as described previously (Hammond et al., 2016). This experimentally determined distance distribution was then compared to the predicted distribution based upon the calculated ensembles of spin label locations at each labelling site. The experimentally measured distances correlate with the predicted distances derived from the chromoATPase crystal structure (3MWY) (Hauk et al., 2010) to within a few angstroms (Figure 2 – figure supplement 2). This suggests that the structure of the chromoATPase domains is similar in solution to that observed in the crystal structure.

In order to study the orientation of the DBD with respect to the chromoATPase domains a series of labelling sites were selected in these domains (Figure 2AB). A total of sixteen distinct distance measurements were made between different pairwise combinations of these sites. Measurements between locations within the DBD and ATPase lobe 1 gave rise to several well defined measurements where an oscillation is evident in the background corrected signal and a single major distance distribution can be extracted (Figure 2 – figure supplement 3). However, measurements between ATPase lobe 2 and the DBD were in general less well defined, often giving rise to multiple distance distributions with similar probabilities (Figure 2 – figure supplement 4). This is likely to arise from increased dynamics between these domains. Similarly, measurements between chromodomains and the DBD did not provide tight single distance distributions (Figure 2 – figure supplement 5). The most prominent distribution of distances for each pair of labelling sites (Shown in Figure 2 – figure supplement 3-5) was used as a constraint in two separate modelling approaches. In the first, the distribution was used to align the centres of the modelled distribution of the MTSL label using Xplor-NIH (Schwieters et al., 2003) (Figure 2C). In the second, TagDock (Smith et al., 2013) was used to fit the distribution of experimentally determined distances to the distribution of modelled spin label locations (Figure 2D). After 500 runs the RMS deviation of the solutions for the Xplor and TagDock based solutions were 2.8 and 1.6Å respectively. In addition, the average of the ensemble solutions obtained using the two approaches has an RMS deviation of 2.4Å. The final averaged solution obtained by TagDock based modelling (Figure 2 – figure supplement 6) was used to compare experimentally measured distances from those predicted by the model (Figure 2 – figure supplement 6). Most of the distances derived from measurements fit the model with a few angstroms deviation. The PELDOR-derived model is compatible with the volume envelopes obtained from the SAXS data (Figure 2 – figure supplement 7A). While the SAXS pattern computed by program CRYSOL (Svergun et al., 1995) from the atomistic EPR-based model of Chd1 yields a poor fit to the experimental scattering data from the full length protein with discrepancy χ^2^= 11.5 (Figure 2 – figure supplement 7B). This misfit is to be expected given that significant parts of the structure (174 and 36 residues at the N‐ and C-termini, respectively, and a 82-residue linker between chromoATPase and DBD domains) are missing in the Chd1 model. The program BUNCH was used to reconstruct probable configurations of these missing portions in the form of dummy residue (DR) chains (Petoukhov and Svergun, 2005). The addition of the missing loops significantly improved the agreement between the experimental and the calculated data yielding χ^2^ = 2.55 (Figure 2 – figure supplement 7B). The radius of gyration for the ensemble of solutions that fit the experimental scattering has a tighter distribution suggesting these regions are not entirely disordered.

**Figure 2.**
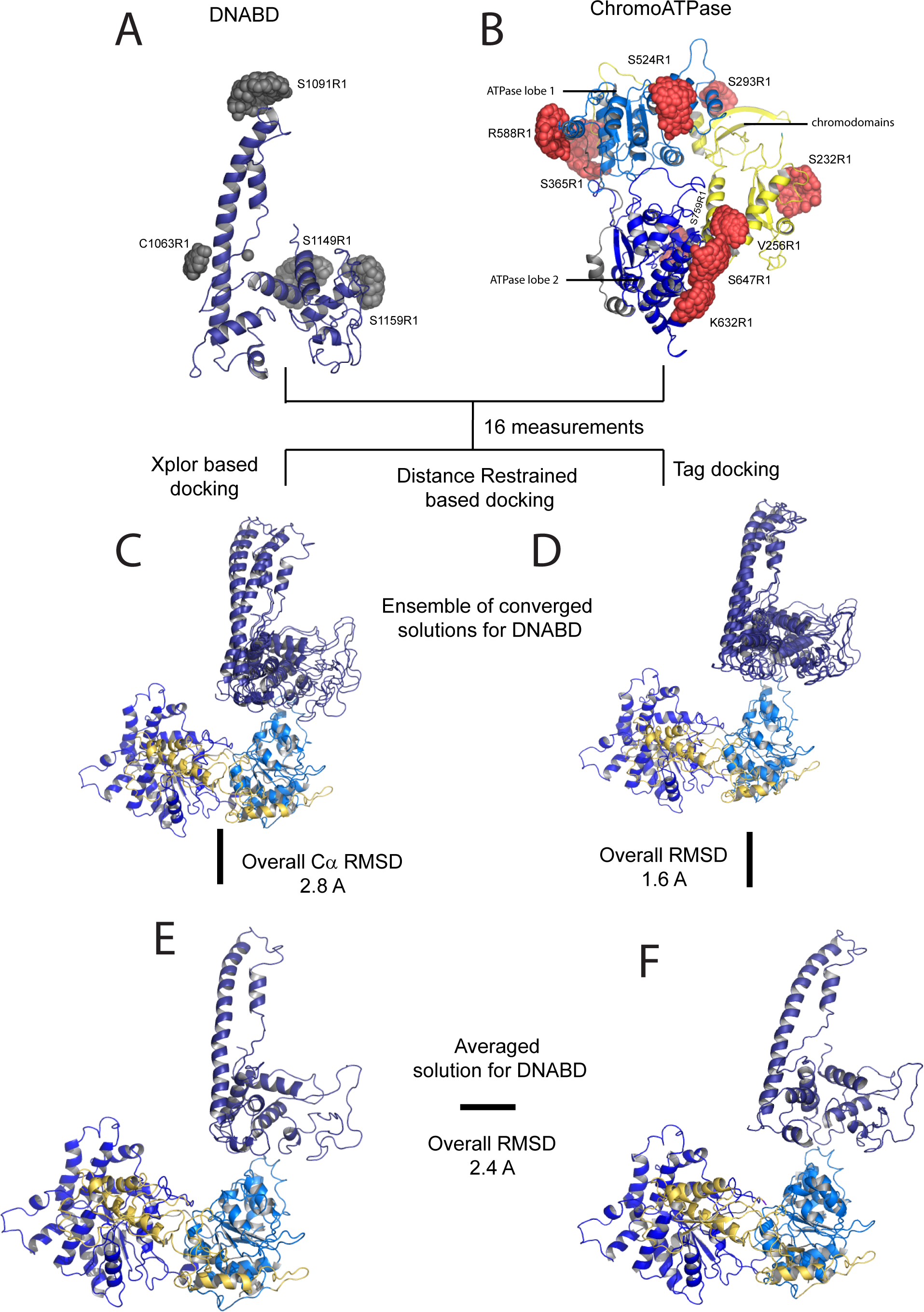
A model for the solution structure of Chd1 based on pulsed EPR measurements. **A**, Structure of DNA binding domain and **B** chromoATPase domains of Chd1 shown in cartoon representation. The ensemble nitroxide atom distributions correspond to different molecular dynamics simulated conformers of the MTSSL spin label and shown as grey spheres. Chromo domain in yellow, ATPase lobe1 in marine, ATPase lobe 2 in blue and the linker region NegC in grey. **C**, Converged solutions of the DNA binding domain orientation relative to the chromo helicase domain determined by rigid body docking using sparse PELDOR data as distance restraints in Xplor-methods. The overall Cα RMSD of the converged structures is indicated. **D**, An alternative approach using the Tagdocking method was amended for rigid body docking of the DNA binding domain on to the chromo helicase. The Cα RMSD of the converged structures is indicated. **E,F** Final averaged structure and the relative Cα RMSD between the structure obtained with two methods are shown in cartoon representation.

### The N-terminus of Chd1 contributes to the interface with the DBD

A striking feature of the model derived for the solution structure of Chd1 is that there is not a significant interaction interface between the DBD and the chromoATPase domains (Figure 2E, F). However, our PELDOR distance measurements clearly show that the DBD is generally constrained with respect to the chromoATPase region. The best explanation for this is that one or more of the regions of Chd1 for which there is no structural information interacts with the DBD and constrain its position. As a means of identifying putative regions that may interact with the DBD we performed chemical crosslinking coupled with mass spectrometry using the amine-reactive crosslinker BS3G (Leitner et al., 2016). BS3G has a length of 11.4Å and so it crosslinks regions of the protein that are relatively close in space. Consistent with this, we identify a number of crosslinks within regions of Chd1 known to be close to one another based on existing structural data, such as between the ATPase lobes and the chromodomains (Figure 3A). Interestingly, several crosslinks between the N-terminal region of Chd1 and the DBD are also identified (Figure 3A), suggesting these two regions are close to one another in the intact protein.

**Figure 3.**
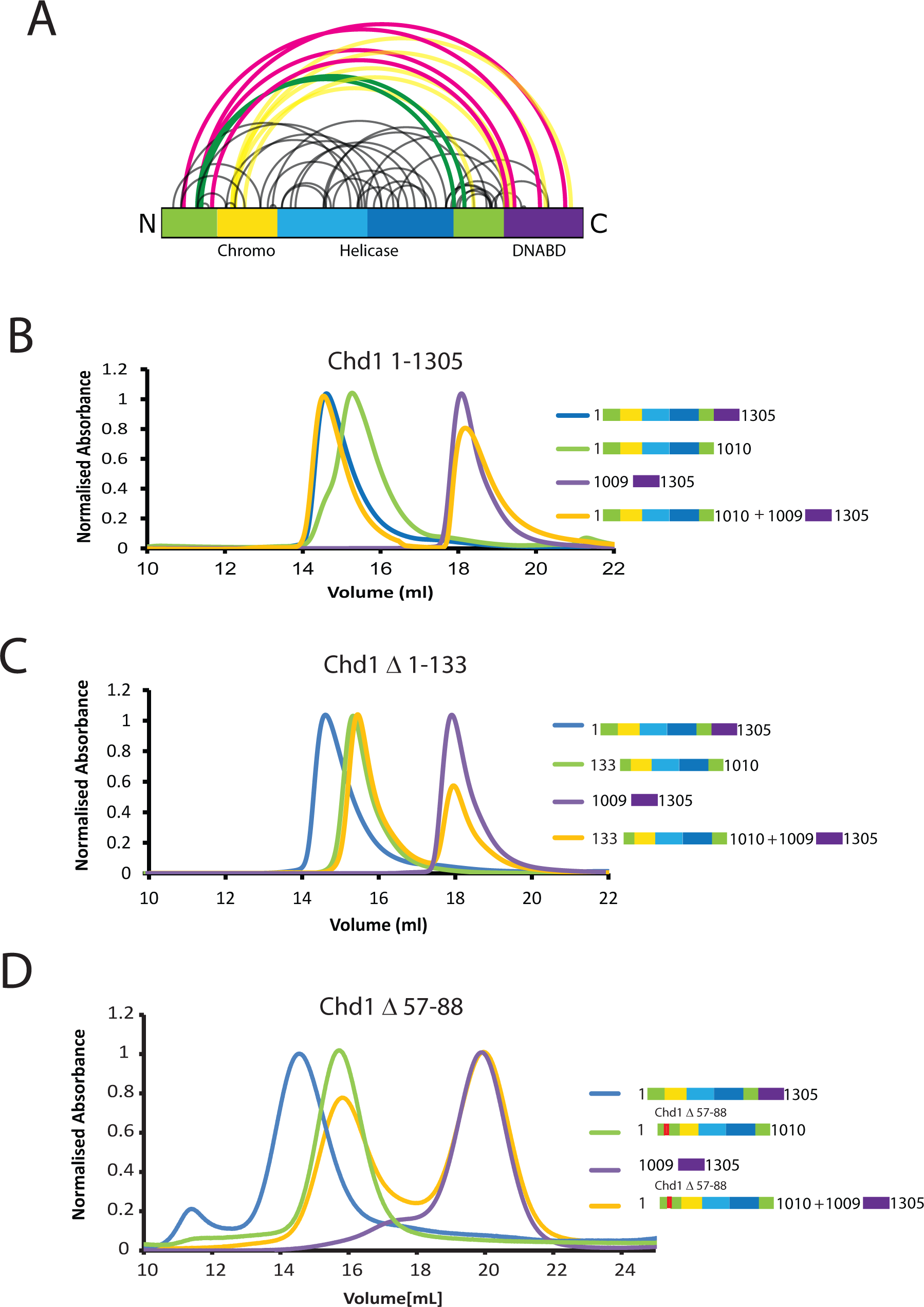
The N-terminus of Chd1 mediates interactions with the DNA binding domain. **A**, Sites covalently linked by the crosslinker BS^3^ were identified by mass spectrometry and are represented graphically using a plot generated with Xvis (Grimm et al., 2015). Thick red and green lines indicates crosslinks between the N-terminal and the C-terminal regions. **B**, Size exclusion chromatography (SEC) elution profiles of selected Chd1 fragments. Normalised elution profile of DBD (1009-1305) in purple, chromo-helicase with intact N-terminal region (1-1010) in green, Chd1 (1-1305) in blue. The elution profile for a 3:1 mixture of the DBD with the chromoATPase (1-1010+1009-1305) is shown in orange. The chromoATPase elutes at low volume consistent with the formation of a complex between the N‐ and C-terminal fragments. **C**, Normalised elution profile of Chromo-helicase with missing the N-terminal 133 amino acids is coloured green. Other profiles are similar to as described in Figure 3A. Loss of the N-terminal 133 amino acids prevents association with the C-terminal DBD. **D**, Similar SEC experiment performed using a Chd1 N-terminal fragments that includes the internal deletion Δ57-88. This also prevents association between the two halves of the protein.

Further evidence supporting an interaction between the DBD and the N-terminal region of the protein comes from gel-filtration experiments. We observe that the isolated recombinant DBD is still able to form an intermolecular complex with a truncated form of Chd1 (residues 1-1010) lacking the DBD (Figure 3B). Furthermore, when the N-terminal 133 amino acids are removed from Chd1 this complex with the DBD is no longer formed (Figure 3C). This suggests that contacts between the N-terminus of Chd1 directly contribute to the association with the DBD. To gain further insight into this we compared the N-terminal sequences of Chd1 proteins across different yeast species (Figure 4A). There is considerable conservation within the 133 amino acids N-terminal to the chromodomains and this includes a tract of acidic amino acids (57-88) followed by a positively charged region extending from 90 to 120. Indeed, when we delete just residues 57–88 the interaction with the DBD is also lost (Figure 3D). Thus, this acidic patch appears to play an important role in mediating interactions with the DBD.

**Figure 4.**
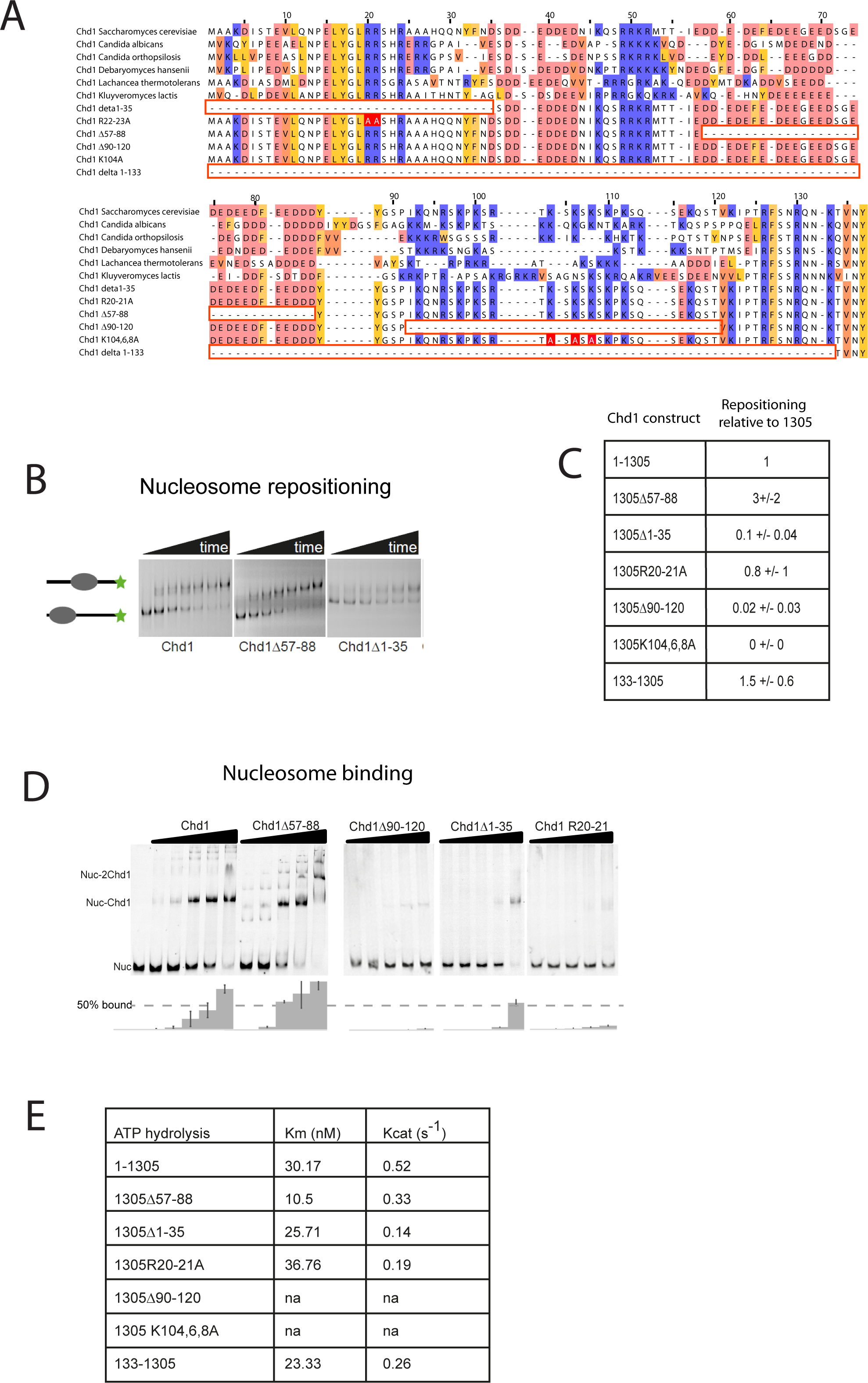
– Mutations to the Chd1 N-terminus have positive and negative effects on activity. (A) Alignment of Chd1 proteins from the indicated yeast species indicates that sequences in the N-terminal region are conserved. Below this mutations to the N-terminal region characterised in this study are indicated. Numbering is to the *Saccharomyces cerevisiae* sequence. (B) 1pmole of nucleosomes assembled on a DNA fragment consisting of the 601 nucleosome positioning sequence flanked by 47bp linker DNA on one side were incubated at 30°C with 20fmoles enzyme, aliquots of the reaction were stopped at 4, 8, 16, 32, 64, and 128minutes. Repositioning of the nucleosome to the DNA centre show an increase in activity for the Δ57-88 and a decrease for the Δ1-35 mutation. (C) Initial reaction rates relative to wild type were determined from a non-linear fit, and are presented as mean +/− standard deviation for N=5 (Δ57-88), 2 (Δ1-35,Δ90-120, K104,6,8A), 4 (R21,22A), and 3 (133-1305). (D) 20nM nucleosomes were incubated with increasing concentrations of enzyme (80, 160, 230, 320, and 460nM in the left panel; 40, 80, 160, and 320nM in the right panel). Representative gel images are shown with bar graphs below showing the mean +/− standard deviation of percent bound from triplicate experiments. (E) 80nM enzyme was reacted with increasing amounts of nucleosome (10, 20, 40, 80, and 120nM) and phosphate release from ATP hydrolysis was monitored by fluorescence intensity. Non-linear regression of triplicate experiments was used to define Km as the nucleosome concentration at half maximal reaction rate and kcat as the enzyme turnover at maximum rate.

### The N-terminus of Chd1 positively and negatively regulates activity

The orientation of the DBD and the chromoATPase domains may influence the ability of Chd1 to productively engage with nucleosomes and therefore influence nucleosome remodelling activity. To assess the contribution of the N-terminal regions to Chd1 activity, mutant Chd1 proteins were expressed in which these regions were mutated. In addition to deletion of the conserved acidic region amino acids 57-88, an adjacent conserved basic region amino acids 90-120 and the entire N-terminus were also deleted. The ability of these proteins to reposition nucleosomes initially located near the ends of DNA fragments towards the centre of the fragments was assessed. Deletion of the acidic patch was observed to increase nucleosome sliding 2.5-fold, while deletion of the basic region reduced activity over 10-fold (Figure 4B,C). The ability of alterations within the N-terminus to either increase or reduce activity was born out by additional mutations. Deletion of the entire N-terminal 133 amino acids increased activity 1.5-fold, while deletion of the extreme N-terminal 35 residues reduced activity 4-fold (Figure 4C). Point mutation of conserved arginine residues, 20 and 21 to alanine within this region also reduced activity (Figure 4C). Triple mutation of three conserved lysine residues within the basic region resulted in a greater than 10-fold reduction in activity (Figure 4C).

These mutants also affected ATPase activity, with reduced or increased activity generally correlating with nucleosome sliding activity (Figure 4E). Two mutants had such low activity that kinetic parameters could not be calculated. Amongst the others, Km and Kcat were affected. Reductions in Kcat were observed for the inhibitory mutations indicating functional interplay with the ATPase domain. The increased activity of the Chd1 Δ 57-88 arose largely as a result of a reduced Km (Figure 4E). This suggests that nucleosome binding was affected, which was studied using gel shift assays. Chd1 Δ 57-88 was found to form complexes with nucleosomes more effectively than intact Chd1 (Figure 4D). The Chd1 Δ 90-120, Δ 1-35 and R20-21A mutations that strongly reduced activity also bound nucleosomes weakly (Figure 4D). As both the Chd1 Δ 57-88 and Δ 1-133 deletions that increase activity reduce association with the DNA binding domain we speculated that a more flexible linkage allows for increased activity while inactivating mutations may orient these domains less favourably for full activity.

The enhancement of activity observed in Chd1 Δ 57-88 is reminiscent of activating mutations observed in other chromatin remodelling ATPases (Clapier and Cairns, 2012; Hauk et al., 2010). To investigate the consequences of this change in activity *in vivo*, full length Chd1 or Chd1 Δ 57-88 were integrated into the *CHD1* locus of an *Isw1ΔChd1Δ* mutant strain. In this strain positioning of coding region nucleosomes is severely compromised and is partially restored when full length Chd1 is reintegrated (Figure 5). When Chd1 Δ 57-88 is reintegrated, a nucleosomal oscillation is restored with slightly greater amplitude than the wild type. As the amplitude of the oscillation is dependent on Chd1, a greater amplitude is consistent with the increased activity observed *in vitro*. In addition, the maximal density of nucleosome dyads is offset downstream by 4 base pairs at nucleosome +3 and increments of approximately 4 base pairs at subsequent downstream nucleosomes (Figure 5 – figure supplement 1). This is consistent with increased spacing between coding region nucleosomes from 16 to 20 base pairs with alignment to the transcriptional start site (TSS) retained. Although this is a relatively small difference, nucleosome spacing is distinct in different yeast species and determined by *trans* acting factors (Hughes et al., 2012; Hughes and Rando, 2015; Tsankov et al., 2010). The observation that Chd1 Δ 57-88 directs increased inter-nucleosome spacing could result if the deletion affects the “measurement” of linker length by the enzyme. Alternatively, it could arise from the increased activity of this protein. To test whether the total amount of Chd1 activity affected inter-nucleosome spacing full length Chd1 was introduced on a higher copy number plasmid, resulting in a >3 – fold increase in expression as assessed by western blotting (Figure 5 – figure supplement 3). The alignment of nucleosomal reads to the genome from this strain indicates a slightly greater increase in nucleosome spacing, approximately 6 base pairs (Figure 5 – figure supplement 1,2). Introduction of Chd1 Δ 57-88 on the high copy pRS423 plasmid caused a further increase in spacing to approximately 26 base pairs (Figure 5 – figure supplement 1,3). These observations are consistent with coding region internucleosome spacing being influenced by the level of Chd1 activity present.

**Figure 5.**
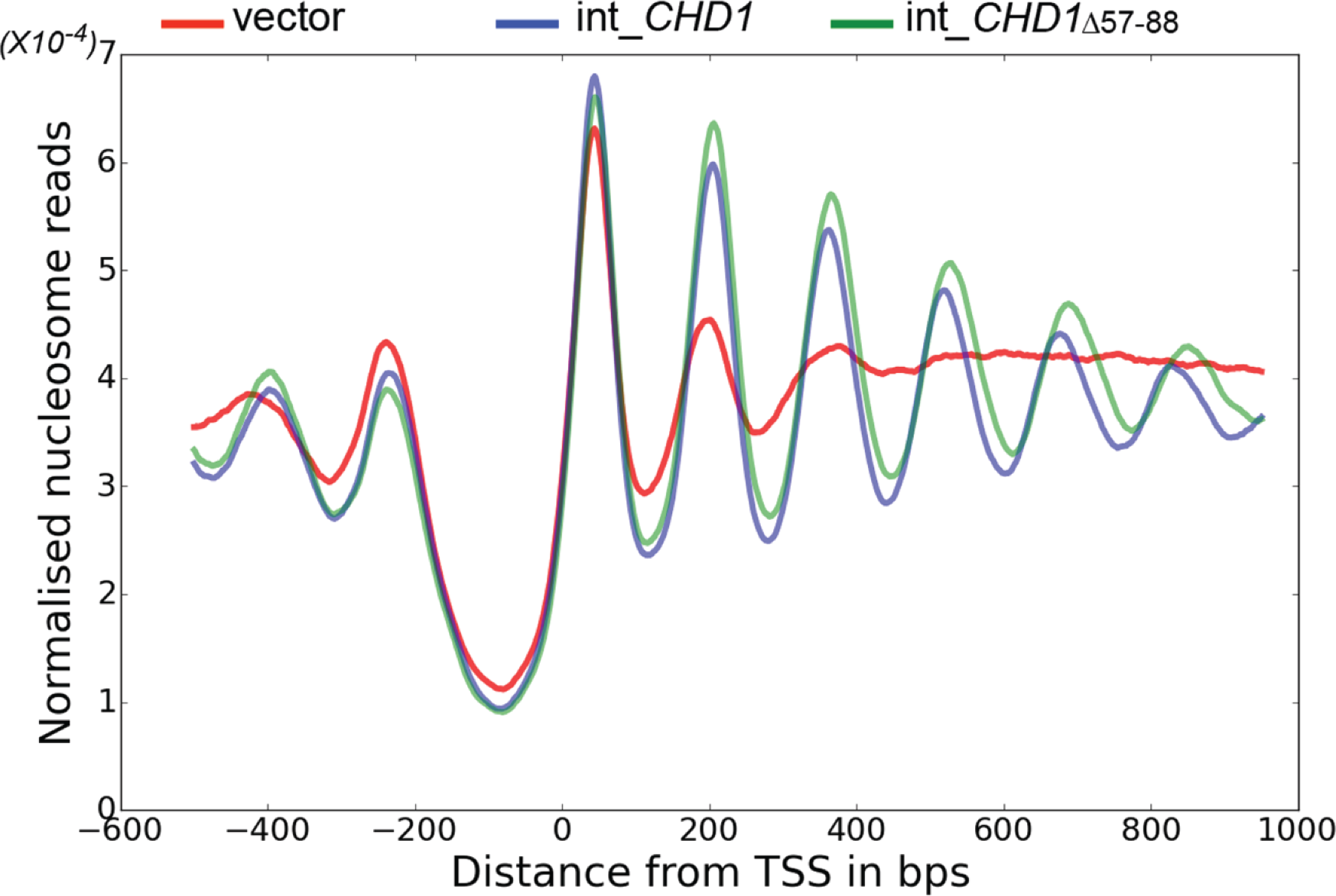
– Chd1Δ57-88 exhibits increased internucleosome spacing in vivo. MNase-Seq was carried out on chd1Δisw1Δtrain transformed with *CHD1* and *CHD1Δc*Δintegrated at *CHD1* lociΔ to obtain genome-wide nucleosome occupancy profiles. TSS-aligned nucleosome occupancy profiles are plotted and show restoration of nucleosome organisation with both Chd1 proteins, but with a downstream shift in the locations of nucleosomes organised by the *CHD15Δ7-88* mutant.

### Interaction of Chd1 with nucleosomes in the apo state

We next sought to study the interaction of Chd1 when engaged with nucleosomes. Cryo-EM was adopted to achieve this and we initially studied the interaction of Chd1 1-1305 with nucleosomes with symmetrical 25-bp linkers using this approach. Under conditions favouring a 1:1 Chd1:nucleosome complex as assessed by native gel electrophoresis, a monodisperse distribution of particles was observed (Figure 6A). Particles picked from micrographs could be assigned to 2D classes, some of which indicated the presence of an attachment adjacent to the nucleosome.

**Figure 6.**
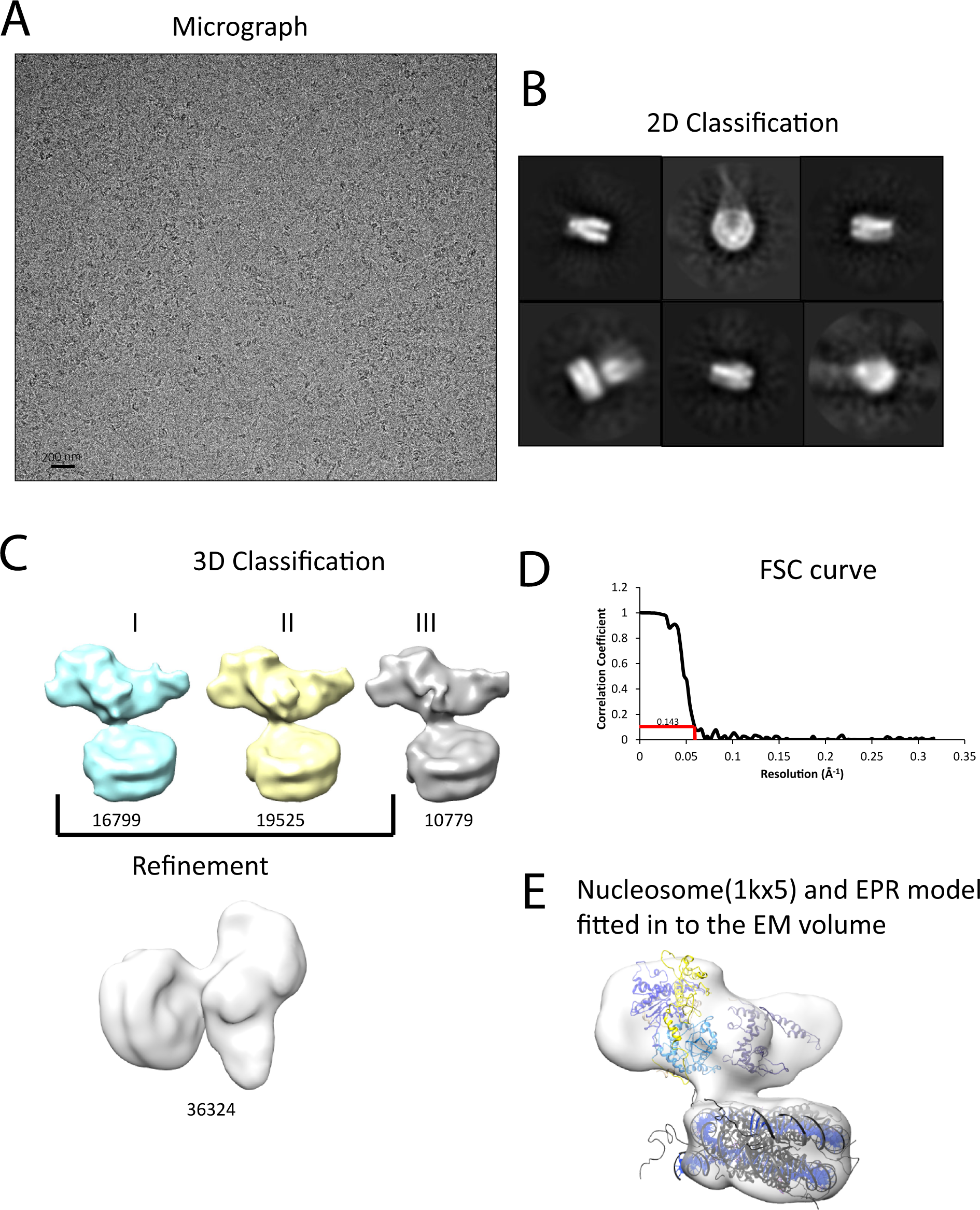
Interaction of Chd1 with nucleosomes in the APO state. **A**, Representative micrograph of frozen hydrated nucleosome-Chd1 apo complex. **B**, Two-dimensional class averages of the CTF corrected auto picked particles is shown. **C**, Volume maps are drawn for the 3 classes obtained from the 3D classification of the particles and final refined volume of the set of 3D classified particles from merging two similar 3D classes as indicated. **D**, Fourier-Shell correlation after gold-standard refinement and conservative resolution estimate at 0.143 correlation. **E**, Electron density map obtained for nucleosome-Chd1 apo complex is shown in semi-transparent surface. The constructed Chd1 solution structure and the nucleosome (1KX5) are docked into the volume.

Following 3D classification and refinement a volume was obtained with a resolution of approximately 20A (Figure 6C,D). This volume is made up of separate areas into which our in solution model for Chd1 and a mononucleosome could be fitted (Figure 6E). A notable feature of this fit is that the engagement surface between Chd1 and the mononucleosome is minimal. It is highly unlikely that this involves the engagement of both the nucleic acid interacting region within the ATPase domains and the DNA binding domain itself. The simplest explanation for this mode of engagement would be that Chd1 is bound through interactions between the DNA binding domain and the ATPase domains are not engaged.

### Characterisation of a fully engaged Chd1-nucleosome complex

We noticed that the hyperactive mutants of Chd1, Δ 57-88, Δ 1-133 formed complexes with nucleosomes that migrated slightly faster on native gels (Figure 4D). This suggests a more compact structure. To characterise this further, complexes between nucleosomes bearing an 11-bp asymmetric linker were formed in the presence of the non-hydrolysable ATP analogue ADP-BeF_x_ which has been observed to stimulate binding of other enzymes to nucleosomes (Leonard and Narlikar, 2015; Racki et al., 2009). These complexes were subject to purification by gel filtration chromatography prior to preparation of grids and micrographs collected using a titan krios microscope equipped with a falcon II detector. 2450 movies were collected and from these 280,000 particles picked. 197602 of these could be assigned into 2D classes that were consistent with a nucleosome with a bound attachment (Figure 7 – figure supplement 1). A first round of 3D classification resulted in the identification of four major classes three of which were similar. The particles from these classes were subjected to 3D auto refine. Subsequently, particle wise movie correction and particle re-centering was carried out and followed by a second round of 3D classification. Solvent masking was applied prior to the final refinement which resulted in a final volume with a resolution of 15Å (Figure 7 – figure supplement 1).

A nucleosome can be docked within the volume and the DNA volume on the side of the nucleosome lacking linker provides a strong reference point to orient the nucleosome (Figure 7; Figure 7 – figure supplement 2). The position of the chromoATPase is also unambiguous and the protrusion formed by ATPase lobe 2 and the chromodomains allows the domains to be oriented (Figure 7; Figure 7 – figure supplement 2). Docking of the DNA binding domain was more difficult most likely as it and the linker DNA it associates with can occupy different conformations. While the volume occupied by the DNA binding domain must be in the general vicinity of the 11-bp linker, solvent masking was required to reveal the trajectory of the linker DNA. The DNA fragment within the co-crystal structure of the SANT-SLIDE domain bound to DNA (Sharma et al., 2011) could then be uniquely oriented with linker DNA on this trajectory.

When the alignment of Chd1 DNA binding and ATPase domains is compared between the solution structure determined by PELDOR and SAXS (Figure 1,2) to that in the nucleosome engaged structure (Figure 7) it is clear that a major change in the orientation of the DNA binding domain with respect to ATPase lobe 1 takes place (Figure 7 – figure supplement 3A). If Chd1 in the conformation observed in solution is docked onto a nucleosome using the location of the chromoATPase domains in the engaged state as a reference point, steric clashes with the histone octamer indicate this is not possible (Figure 7 – figure supplement 3B). Conversely, docking the solution structure using the DNA binding domain as a reference point results in a configuration more similar to that observed in Figure 6 (Figure 7 – figure supplement 3C).

**Figure 7.**
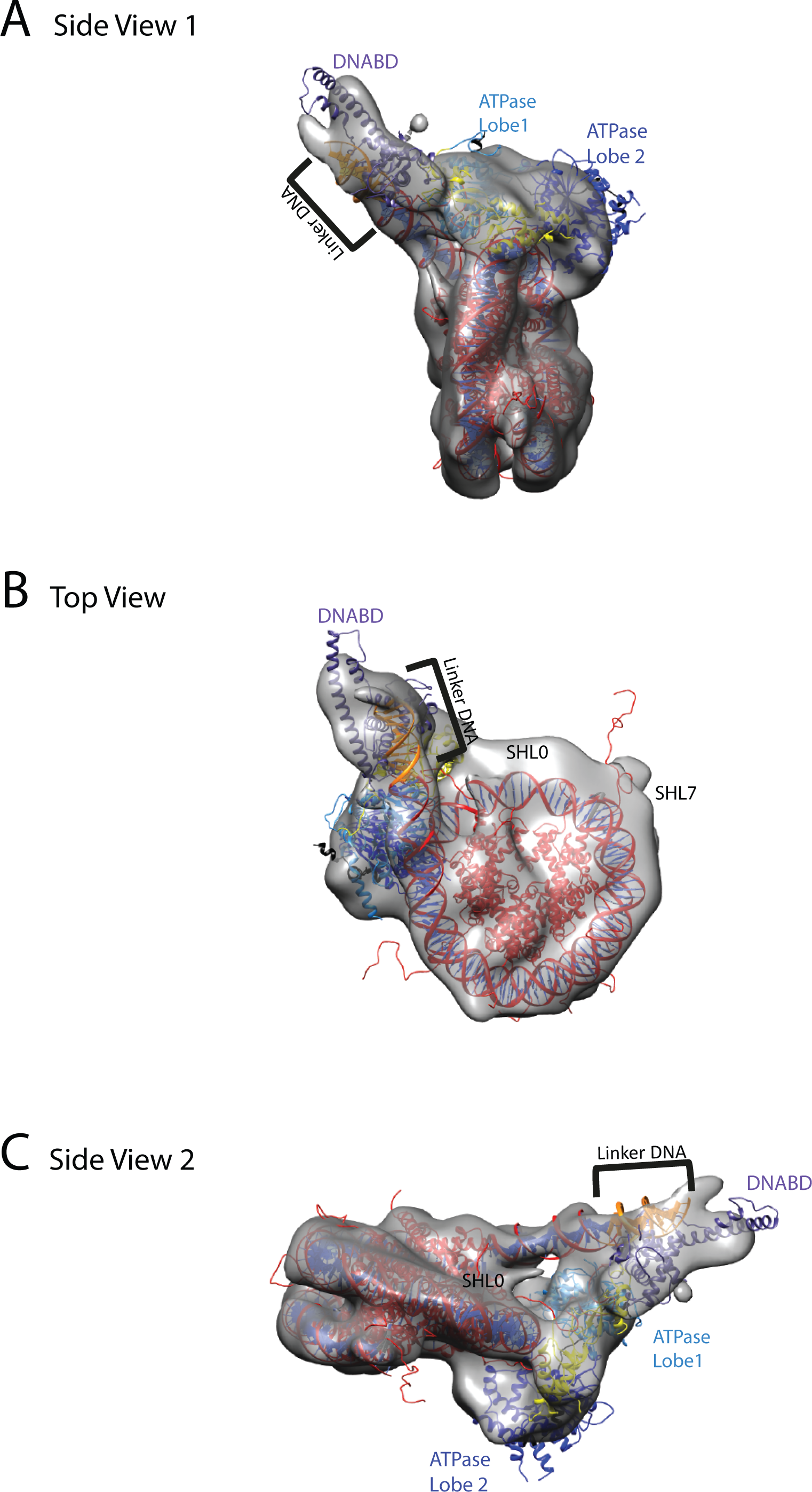
Chd1 bound to nucleosomes via its DNA binding and ATPase domains. **A**, Side view 1 of the electron density map shown in semi-transparent grey surface with docked nucleosome (red, 1KX5) and the Chd1, chromoATPase (3MWY) and DNA binding domain (3TED) crystal structures shown in cartoon representation. The various domains of Chd1 are labelled. ATPase lobe1 in marine, ATPase lobe2 in blue, chromo domain in yellow and the DNA binding domain in deep blue. The 11-bp DNA linker region defined in the electron density map is coloured orange and indicated. **B**, Top view of the nucleosome bound Chd1 complex. The dyad axis of the nucleosome is labelled as SHL0 (super helical location 0) and edge of the nucleosome is indicated as SHL7. **C**, Side view 2 of the nucleosome-Chd1 complex.

### One helical turn of nucleosomal DNA dissociates from the octamer in the engaged complex

A prominent feature of the structural model is that the nucleosomal DNA adjacent to the linker is unravelled from the octamer surface. This is evident as when intact nucleosomal DNA is modelled into the volume it protrudes into a region of missing density (Figure 7 – figure supplement 3). In addition, a channel of volume contiguous with the prominent nucleosomal DNA at SHL6 emerges from the nucleosome on the side bearing extranucleosomal DNA (Figure 7 – figure supplement 3).

To characterise this further we made use of a nano-positioning approach that measures fluorescence resonance energy transfer (FRET) between dye molecules introduced at specific locations (Muschielok et al., 2008). Fluorescent dyes were introduced to DNA derived from the 601 nucleosome positioning sequence on either the forward (F) or reverse (R) strand at the indicated positions relative to the nucleosomal dyad axis of symmetry (Figure 8A; Figure 8 – figure supplement 1). Nucleosomes were assembled onto fragments with a 6 base pair extension to one side of the nucleosome and a 47 base pair linker on the other side 5’ labelled with biotin to provide a means of coupling to a streptavidin coated slide. When the FRET efficiency was measured between Alexa647 at F-71 and Tamra at F+14 the mean FRET efficiency was 0.8 consistent with nucleosomal wrapping bringing these two sites into close proximity. When Chd1 1-1305 was added to these nucleosomes no change in FRET efficiency was observed (Figure 8B). This is consistent with the 6 base linker on this side of the nucleosome being too short to direct binding of the Chd1 DBD. In contrast, when nucleosomal DNA was labelled at F+2 with Tamra and at R-66 with Alexa647, binding of Chd1 1-1305 in presence of AMP-PNP caused a reduction in the mean FRET efficiency from 0.5 to 0.3 (Figure 8C). This is consistent with Chd1 binding causing DNA 9 base pairs from the edge of the nucleosome to dissociate in a fashion similar to that observed with Chd1 Δ57-88 by single particle cryo EM (Figure 7; Figure 7 – figure supplement 2). The dissociation of nucleosomal DNA did not extend deep in the nucleosome as a reporter site at R-60 was unaffected by Chd1 binding (Figure 8D). A site further into the linker, namely R-85 showed a larger change in FRET efficiency from 0.45 to 0.1 upon binding of Chd1 (Figure 8 – figure supplement 2). The larger change in FRET efficiency at this location made it more tractable to assess dynamic changes in FRET quantitatively. This made it possible to show that the reduction in FRET efficiency was more prominent in the presence of Chd1 1-1305 and AMP-PNP than it was in the presence of Chd1 1-1305 alone (Figure 8 – figure supplement 2B). Changes in FRET could be tracked on individual complexes either in the absence (Figure 8 – figure supplement 2C) or the presence of AMP-PNP (Figure 8 – figure supplement 2D). Statistical analysis of several hundred single molecule traces (Figure 8 – figure supplement 2E) allowed to extract kinetic rates for this conformational changes. In the presence of AMP-PNP the unwrapped state was favoured both as a result of an increased rate of transition to the unwrapped state and a reduction to the rate of DNA re-association (Figure 8 – figure supplement 2F). These observations indicate that DNA unwrapping occurs with a non-mutant form of Chd1 similar to that observed with Chd1 Δ57-88 by single particle EM (Figure 7). In addition, unwrapping is favoured in a nucleotide bound state consistent with the distinct configurations of Chd1 observed in the absence (Figure 6) and presence of ADP-BeF_x_ (Figure 7). The dissociation of the outer turn of DNA during the formation of this engaged complex may serve to prime the nucleosome for dynamics driven by ATP-hydrolysis.

**Figure 8.**
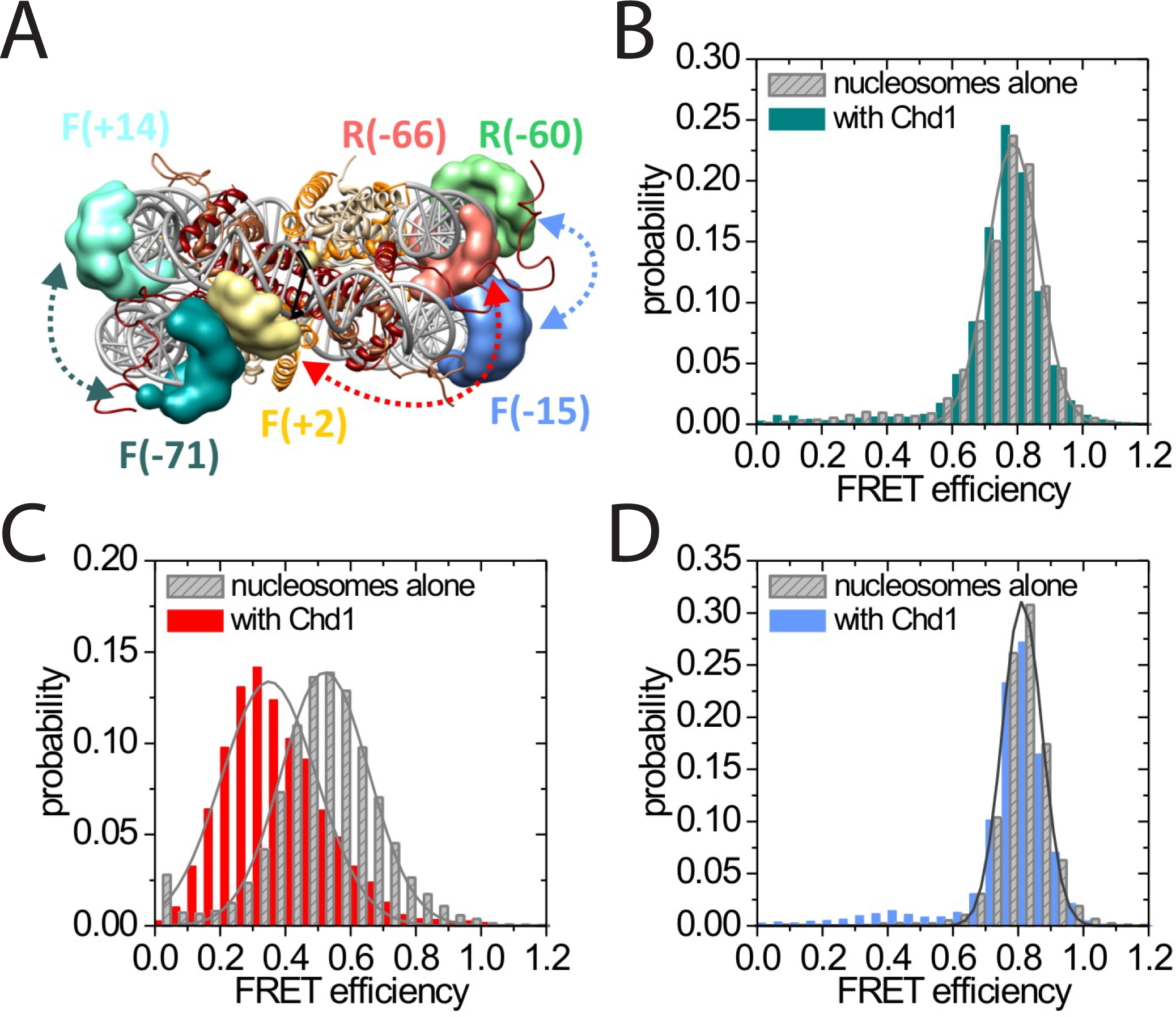
Nucleotide binding affects DNA wrapping in Chd1-nucleosome complexes. Experiments were performed using nucleosomes with dye labels attached to two specific positions alone and in presence of Chd1 1-1305. (**A**) Schematic illustration of the dye positions for the smFRET measurements. (**B-D**) Histograms of measured smFRET efficiencies for nucleosomes alone (grey) and in presence of Chd1 1-1305 and 150μM AMP-PNP (colored) for dyes attached to positions F-71 and to F+14 (B, dark cyan), to F+2-Tamra and to R-66 (C, red) and to F-15 and to R-60-Alexa647 (D, light blue).

## Discussion

In this study we built models for the structure of the remodelling enzyme Chd1, both in free solution and when engaged with nucleosomes. We observe that Chd1 can engage with the nucleosome in different modes (Figure 6, 7). It is likely that Chd1 can interact with nucleosomes in additional conformations as this study is not exhaustive in characterising a full spectrum of complexes that can be formed with different linker DNA lengths and in different stages of ATP hydrolysis. In addition, complexes with two molecules of Chd1 bound to one nucleosome can be formed but have not been characterised here. Because Chd1 can engage with nucleosomes in different ways, it is possible that ensemble approaches that have previously used to characterise the interaction of Chd1 and other remodelling enzymes with nucleosomes may report on a mixture of different states.

In the absence of ATP analogues, Chd1 is bound to the nucleosome through a small interaction surface (Figure 6). The SANT-SLIDE domains of Chd1 and the related domains in ISWI, bind DNA with affinities in the low nanomolar range and are required for efficient interaction with nucleosomes (Grune et al., 2003; Ryan et al., 2011). DNA recognition by Chd1 does not appear to be sequence specific (Sharma et al., 2011). Consistent with this, the related DNA binding domain of Isw1a has been observed to interact with different regions of linker DNA dependent on the nucleosomal substrate used (Yamada et al., 2011). It is possible that interaction of the DNA binding domain with linker DNA represents a first step in the association of Chd1 with chromatin. One dimensional diffusive motion that has been observed for many non-specific DNA binding proteins (Stanford et al., 2000) provides a means for enrichment on longer linker DNA and a means of scanning exposed linker DNA for sites where engagement of the ATPase domains with nucleosomes is also possible.

In Figure 7 we observe a hyperactive mutant of Chd1 bound to a nucleosome bearing an 11-bp linker in the presence of the ground state mimic ADP-BeF_x_. In this case a more compact particle is observed consistent with engagement of both the ATPase and DNA binding domains (Figure 7). As structural data has previously been obtained for the nucleosome, Chd1 chromoATPase domains and SANT-SLIDE DNA binding domain it is possible to orient these major components within the volume obtained (Figure 7). The DNA binding domain is bound close to the edge of the nucleosome interacting with the first 10 base pairs of linker DNA. Previous studies have shown that Chd1 activity is greatly reduced when the SANT and SLIDE domains that comprise the C-terminal DNA binding domain are deleted (Hauk et al., 2010; Ryan et al., 2011). Remarkably fusion of Chd1 to heterologous domains that direct specific interactions with DNA restores activity (McKnight et al., 2011; Patel et al., 2012). Chd1-AraC hybrids are dependent on an *araI* site and the optimal position for this is +8-bp from the nucleosome boundary (McKnight et al., 2011). Chd1-streptavidin fusions are dependent on the introduction of biotinylated nucleotides with optimal location +11-bp from the nucleosome boundary (Patel et al., 2012). Although these fusion proteins will not pack identically to the native SANT and SLIDE domains, these observations indicate that an interaction with the first turn of DNA outside of the nucleosome, as observed in our engaged structure, is functionally important. They also show that nucleosome repositioning is directed towards the bound linker (McKnight et al., 2011; Patel et al., 2012).

The outer turn of nucleosomal DNA is removed from the surface of the histone octamer on the side of the nucleosome with the linker DNA bound to the Chd1 DNA binding domain (Figure 7 – figure supplement 4). This occurs in the absence of ATP-hydrolysis, but may be facilitated by conformational changes promoted by the binding of the transition state mimics ADP-BeF_x_ or AMP-PNP (Figure 8 – figure supplement 2). Consistent with this, the related enzyme Snf2h has also been observed to bind nucleosomes more effectively in the presence of ADP-BeF_x_ (Racki et al., 2009) and ISW2 protects from hydroxyl radical digestion more effectively in the presence of bound nucleotides (Gangaraju et al., 2009). More effective engagement of the ATPase domains may favour the binding of the DBD to linker DNA. This in turn may cause dissociation of nucleosomal DNA in a similar fashion to that observed upon binding of sequence specific DNA binding proteins (Adams and Workman, 1995; Polach and Widom, 1996). The loss of octamer contacts on the bound side of the nucleosome will reduce the energy required to affect dynamic alterations to nucleosomal DNA and may act as a priming step for subsequent ATP-dependent remodelling. This priming step may distinguish the initial ATP dependent changes from subsequent cycles of activity. Consistent with this 7 base pairs of DNA have been observed to be removed from nucleosomes following the action of ISW2 and Isw1b complexes followed by subsequent 3 base pair movements (Deindl et al., 2013).

ATP-dependent repositioning occurs as a result of the action of the chromoATPase domain which engages with nucleosomal DNA, centred 1.5 superhelical turns off the nucleosomal dyad (Figure 7). There are two SHL 1.5 locations either side of the nucleosome. The SHL 1.5 location that is bound is further away on the linear DNA sequence, separated by 90 rather than 60 base pairs from the bound linker. However, it is in closer spatial proximity once DNA has been wrapped around the nucleosome (Figure 7). This arrangement is different to that previously reported for ISW2. In this case the proximal dyad location in the DNA sequence is bound by the ATPase domains (Dang and Bartholomew, 2007). The distinct topography with which the domains of Chd1 engage with nucleosomal DNA raise the possibility that the orientation of DNA translocation is also different to that observed with other enzymes (Saha et al., 2005; Zofall et al., 2006). High resolution structures enable the strand specificity of nucleic acid contacts to be assigned (Sengoku et al., 2006). Unfortunately, this structure is not at a resolution where this assignment can be made, so the directionality of translocation cannot be resolved. Similarly, the limited resolution does not enable us to comment on rearrangements that are anticipated to occur between the chromo and ATPase domains and within the ATPase domains during engagement with DNA (Hauk and Bowman, 2011).

It is clear that there is a large change in the orientation of the DNA binding and chromoATPase domains upon engagement with nucleosomes (Figure 7 – figure supplement 3A). A large scale nucleotide dependent change in conformation of the related enzyme SNF2h has been reported previously (Leonard and Narlikar, 2015). This involves repositioning of the SNF2h SANT-SLIDE DNA binding domain from a site that generates a FRET signal with a site +20-bp from the nucleosome edge to one that generates FRET at a site at -25 internal to a nucleosome. It is possible that this could involve redistribution of the SNF2h SANT SLIDE domain from positions distributed along the linker to high occupancy at the nucleosome boundary. Additional evidence for a conformational change upon binding to DNA and nucleosomes has been obtained by studying changes in protease digestion of Drosophila ISWI following binding to DNA (Mueller-Planitz et al., 2013).

If the predominant conformation observed in solution is docked onto a nucleosome using the engaged chromoATPase domains (Figure 7) as a reference point, steric clashes occur with the histone octamer (Figure 7 – figure supplement 3B). However, if the solution structure is docked using the DNA binding domain from Figure 7, a conformation related to that of the apo enzyme (Figure 6) is observed (Figure 7 – figure supplement 3C). It is likely that there is some interplay between these states. Many of the solution measurements include minor subpopulations that indicate conformational flexibility, for example between the DBD and ATPase lobe 2 (Figure 2 – figure supplement 5). These minor populations do not provide an obvious fit to the fully engaged conformation. Instead, ATPase lobe 2 may be more mobile perhaps reflecting the different conformations that have been observed in crystal structures (Durr et al., 2005; Thoma et al., 2005; Xia et al., 2016). Nonetheless, the major species observed in solution is not capable of binding nucleosomes in the fully engaged conformation which we assume to be active. As a result the native protein could be considered to be auto inhibited. Consistent with this, both within Chd1 (Figure 4; (Hauk et al., 2010)) and other remodelling ATPases (Clapier and Cairns, 2012), mutations have been reported that increase activity. Here we show that mutations to the N-terminus that increase activity, reduce interactions between the DBD and the chromoATPase (Figure 4B,C). This suggests that increased flexibility in the orientation of the chromoATPase and DBD increases the opportunity for productive engagement with nucleosomes. As a result, regulation of accessory domain orientation to enable engagement with the nucleosomal substrate represents a potent means of regulating the activity of remodelling ATPases. This may be important as the combined abundance of chromatin remodelling enzymes (Flaus and Owen-Hughes, 2011) could present a load on nuclear ATP levels. Regulation allowing increased activity when chromatin organisation is required, for example following transcription or DNA replication would reduce the energy expended organising chromatin.

Chd1 plays a major role in the maintenance of even nucleosome spacing within coding regions (Gkikopoulos et al., 2011). To achieve this, the activity of the enzyme is also anticipated to be influenced by the length of DNA between adjacent nucleosomes. Within the ISWI related ACF complex, the duration of pauses prior to repositioning is shortened when linker lengths are increased in the range 40 to 60-bp (Hwang et al., 2014). An activation mutation termed AutoN (Clapier and Cairns, 2012), N-terminal to the ATPase domains, is found to impede DNA length sensing (Hwang et al., 2014). This may function in a similar way to the hyperactive Chd1 Δ 57-88 mutant, enabling engagement with nucleosomes in an active conformation independent of linker DNA length. However, there may also be important differences in the way in which the SANT-SLIDE domains of ISWI and Chd1 related proteins act. The structures of these domains do differ significantly in ISWI and Chd1 (Grune et al., 2003; Ryan et al., 2011). In addition, yeast Chd1 acts to organise nucleosomes with relatively short linker lengths of 16 base pairs within yeast coding regions (Gkikopoulos et al., 2011; Ocampo et al., 2016). In this respect engagement with the first turn of extranucleosomal DNA as we observe (Figure 7) provides a means of achieving activity within close packed nucleosome arrays. At the same time if Chd1 relocates nucleosomes to less than 20 base pairs from adjacent nucleosomes steric clashes with the SANT-SLIDE bound linker would be anticipated to get progressively more serious and reduce enzyme activity. The inhibition of repositioning when linker length falls below a low limit is sufficient to drive the even spacing of nucleosomes via a bidirectional process (Blosser et al., 2009). Studies of additional intact remodelling enzymes with nucleosomes will be required to determine which features of Chd1 are conserved.

## Methods

### Cloning, protein expression and purification

ScChd1 C-terminal and N-terminal truncations were made from the full length clone described in Ryan et al, using an inverse PCR strategy (Ryan et al., 2011). Site directed mutagenesis was used to introduce cysteine residues at strategic locations on ScChd1 1-1305ΔC. All proteins were expressed in Rosetta2 (DE3) pLysS *Escherichia Coli* cells at 20° C in auto-induction media and the purification of the protein was carried out as described in Ryan et al. After purification, the GST-tag was cleaved with Precission protease and the cleaved proteins were subjected to size exclusion chromatography using Superdex S200 10/300 GL columns (GE Healthcare).

### Assembly of recombinant nucleosomes

DNA fragments including the 601 nucleosome positioning sequence was PCR amplified from pGEM-601 template (Lowary and Widom, 1998). DNA products, pooled from 96-well PCR plates were concentrated by precipitation with ethanol, re-dissolved in 1mM EDTA and 5mM Tris-HCl pH 8.0, and purified by anion exchange over a Source 15Q column using a linear NaCl gradient from 200mM to 2M NaCl. The fractions containing the DNA were pooled and concentrated by ethanol precipitation. Expression and purification of *Xenopus laevis* histones and reconstitution of histone octamer were carried out as described previously (Luger et al., 1999). Nucleosomes were assembled by salt dialysis as described previously (Luger et al., 1999). For Single molecule FRET experiments DNA was also generated by PCR, then purified by size exclusion chromatography on a Superose 6 column. Nucleosomes were assembled by salt gradient dialysis using recombinant human histones.

### Small-Angle X-ray Scattering (SAXS)

The SAXS measurements for the five ScChd1 constructs (1-1305ΔC, ΔN133-1305ΔC, 1-1010 ΔC, ΔN133-1010 and ΔN1009-1274ΔC) were performed with a fixed X-ray wavelength of 1.54Å at the EMBL BioSAXS beamline X33 at the DORIS storage ring, DESY (Hamburg, Germany). A photon counting PILATUS 1M detector placed at a distance of 2.7m was used to record the scattered X-rays. A bovine serum albumin solution at 4.5 mg/ml in 50mM Hepes pH 7.5 was used for calibrating the molecular mass. For each sample, scattering data were measured at 10°C, first at high concentration and then for serially diluted samples to check for any concentration dependence of the scattering profiles. Scattering data for solvent blanks were collected for each samples to account for buffer contribution.

Data were processed using Atsas suite, PRIMUS (Konarev et al., 2003). The buffer subtracted data were extrapolated to infinite dilution using Guinier plot. The zero concentration intensity I(0) at low resolution *q*=0 and the radii of gyration (*R*g) were determined from the Guinier approximation. The data were then merged to obtain interference free scattering curves and used in further analysis. Maximum complex dimensions *D*^max^ and the interatomic distance distribution functions *P*(r) were calculated using GNOM. The excluded (Porod) particle volumes were calculated using PRIMUS. The program CRYSOL (Svergun et al., 1995) was used to calculate the theoretical X-ray scattering profile from the EPR based model and the known crystal structure fragments.

*Ab-initio* reconstruction of scattering data was performed using GASBOR (Svergun et al., 2001) which describes it as a chain of dummy residues, with structure minimisation being carried out independently against the composite, merged scattering curve and their pair distribution function *P*(r). Resulting models were then aligned using SUBCOMB. The most typical reconstructions were averaged and filtered using the DAMAVER and DAMFILT packages. The criterion for including the models in the averaging process was based on their normalised spatial discrepancy (NSD) ≤ 1. For unstructured loops dummy residues were fitted using the programme BUNCH (Petoukhov and Svergun, 2005). Theortical scattering curves were generated from the model using CRYSOL (Svergun et al., 1995). SAXS data and models will be released upon acceptance at SASBDB (www.sasbdb.org).

### Spin labelling of ScChd1, PELDOR measurements and modelling

MTSL was conjugated to introduced cysteines immediately following size exclusion purification as described in Hammond et al (Hammond et al., 2016). Excess unreacted labels were removed from the sample by dialysis. PELDOR experiments were conducted at Q-band (34GHz) operating on a Bruker ELEXSYS E580 spectrometer with a probe head supporting a cylindrical resonator ER 5106QT-2w and a Bruker 400U second microwave source unit as described previously (Hammond et al., 2016). All measurements reported here were made at 50K. Data analysis was carried out using the DeerAnalysis 2013 package (Jeschke and Polyhach, 2007). The dipolar coupling evolution data were first corrected to remove background decay. Tikhonov regularisation was then applied to obtain the most appropriate distance distributions from each dataset.

Crystal structures of chromo helicase (PDB Code: 3MWY)(Hauk et al., 2010) and DNA binding domain (PDB Code: 2XB0)(Ryan et al., 2011) proteins were docked together by performing distance restrained rigid body refinement in XPLOR-NIH (Schwieters et al., 2003). For each structure, R1 spin labels were added and the distribution simulated using molecular dynamics and energy minimisation at specific residue positions, using Xplor-NIH MTSSL parameter and topology files. Experimental modal distances were applied as restraints utilising the distance averaging procedures built into the software. Fragments were docked as rigid bodies using Powell minimisation. A set of 1000 docked structures were generated, each randomising the relative orientation of the starting structure. The interaction and the NOE energies of the final docked structures were evaluated and the solutions, that agreed best to the experimental distance distribution, were accepted for the final model.

In addition to the XPLOR-NIH approach, the Tagdock program (Smith et al., 2013) was used to perform distance restrained rigid docking calculations. In brief, 100,000 decoys were generated during the first low resolution docking phase with a random starting orientation. A Boltzmann-weighted MTSSL rotamer library (Smith et al., 2013) was utilised for the respective spin labelled positions. A score for the docked complex was calculated based on the experimental distance restraints. The best scoring 200 structures were then taken to the high resolution refinement stage where finer rotations and translations were applied. The structures that provided improved scores as well as good agreement with the experimental distances were accepted for the final model.

### Chemical crosslinking and MS analysis

Chemical crosslinking experiments on ScChd1 1305 protein was carried out using a BS^3^ crosslinker [Thermo Fisher Scientific]. Crosslinker BS^3^ was prepared at a concentration of 2mM in DMSO. 5μL of the crosslinker was then added to 145μL 20μM ScChd1 1305 enzyme in 40mM HEPES pH 7.8 and 400mM NaCl. The reaction was incubated on ice for 120 minutes and then quenched by the addition of 10 μL 100mM ammonium bicarbonate. Nonspecific crosslinked products (i.e. multimers of ScChd1) were removed by gel filtration on a PC3.2/30 (2.4 mL) Superdex 200 gel filtration column (GE healthcare, UK) that was pre-equilibrated in 100mM ammonium bicarbonate pH 8.0, 350mM NaCl. Fractions containing the samples were pooled together, reduced with 10mM DTT and alkylated with iodoacetamide (20mM). The samples were then sequentially digested first with Trypsin (sequencing grade, Promega UK) overnight at 37 °C and then with GluC, using a protein to enzyme ratio of 20:1 for both digestions. The digested peptides were size exclusion gel filtrated using the Sephadex G-50 resin on a 50% acetonitrile in 100mM ammonium bicarbonate buffer. The fractions were pooled and dried using centrifugal evaporation at 40°C and resuspended in 30 μL of 5% formic acid. The peptide mixture (1 μL) was injected onto a 15 cm EasySpray C18 column (Thermo Fisher) and separated by a linear organic gradient from 2 to 35% buffer B (80% acetonitrile, 0.1% formic acid) over 110 min. Peptide ions were generated by electrospray ionization from the EasySpray source and introduced to a Q Exactive mass spectrometer (Thermo Fisher). Intact peptide ion and fragment ion peaks were extracted from the RAW format into the MGF format using the Proteome Discoverer 1.4 software package, for subsequent analysis. Crosswork algorithm (MassAI software) (Rasmussen et al., 2011) was used to analyse and annotate the crosslinked peptides. The Xvis (Grimm et al., 2015) software was used to represent the annotated crosslinked sites.

### Nucleosome repositioning

Nucleosome repositioning assays were performed in 40mM Tris pH 7.5, 50mM potassium chloride, and 3mM magnesium chloride. Repositioning of 1pmole yeast nucleosomes on Cy5 labelled 0W47 DNA by 20 fmoles of Chd1 enzyme was assessed. Central and edge aligned nucleosomes were separated on a pre-run 6% polyacrylamide gel (49:1 acrylamide: bis-acrylamide) in 0.2X TBE buffer with buffer recirculation at 300V in the cold. Nucleosomes were visualised on Fujifilm FLA-5100 imaging system at 635nm. Percent repositioning was determined using Aida Image Analyser, and data were fit via dynamic curve fit non-linear regression in Sigma Plot. In order to obtain the initial rate of sliding, the derivative of the non-linear fit was solved at t=0.

### Nucleosome binding

*Xenopus laevis* nucleosomes (20nM), reconstituted on Cy3 labelled 0W11 DNA, were bound to titrations of Chd1 enzymes (concentration specified in figure legend) in 50mM Tris pH 7.5, 50mM sodium chloride, and 3mM magnesium chloride supplemented with 100μg/mL BSA. Unbound and bound nucleosomes were separated on a pre-run 6% polyacrylamide gel (49:1 acrylamide: bis-acrylamide) in 0.5X TBE buffer for 1 hour at 150V. The gel shift was scanned on Fujifilm FLA-5100 imaging system at 532nm. The percent of bound nucleosomes was calculated using Aida Image Analyser.

### ATP hydrolysis

ATP hydrolysis measurements were performed in 50uL reaction volumes, containing 3μM phosphate sensor, 200μM Pi-free ATP, and a titration of nucleosome concentrations (specified in figure) in 50mM Tris pH 7.5, 50mM sodium chloride, and 1mM magnesium chloride. Phosphate release was measured as fluorescence intensity by Varian Cary Eclipse with excitation and emission wavelengths of 430 and 460nm, respectively. All reaction components, except for ATP, were added and fluorescence signal was recorded briefly, upon the addition of ATP, the rate of increase in fluorescence was measured. Data were fit via dynamic curve fit, non-linear regression in Sigma Plot, to determine Km, Vmax, and kcat.

### *In-vivo* nucleosome mapping

Yeast culture, nucleosomal DNA preparations and Bioinformatic analysis were carried out as described previously (Gkikopoulos et al., 2011). Strain TOH1482: *chd1::URA3 isw1::HphMX4* was generated form BMA64 (Gkikopoulos et al., 2011). To construct strains with *CHD1*(wt/mutant) integrated at *CHD1* loci, TOH1482 was transformed with *CHD1* constructs (promoter*CHD1-CHD1(*wt/mutant)-3XFLAG-*CHD1* terminator) and selected on 5-FOA for transformats. Strains with increased expression were obtained by transforming TOH1482 strain with the plasmid constructs (pRS423-promoter*CHD1*-*CHD1(*wt/mutant)-3XFLAG-*CHD1* terminator). The *CHD1-* protein levels in each strain was measured by western blot using anti-FLAG antibodies. Triplicate repeats of the genomic datasets are submitted at European nucleotide Archive and are available under study accession number PRJEB15701 (http://www.ebi.ac.uk/ena/data/view/PRJEB15701).

## Cryo-Electron Microscopy (Cryo-EM)

### Sample preparation

The appropriate ratio of ScChd1 to nucleosome for 1:1 complex formation was determined by titration and native PAGE analysis. The complex was then purified by size exclusion over a PC 3.2/30 Superdex 200 column in 20mM Tris, 50mM NaCl. Fractions were analysed by 6% Native PAGE and fractions containing ScChd1-nucleosome complexes pooled. A 4 μL drop of sample was applied to Quantifoil Holey carbon foil (400 mesh R1.2/1.3μm) treated with glow discharge (Quorum technologies). After 15 second incubation at 4 °C, grids were double side blotted for 3.5 s with a blot force of 5 in a FEI cryo-plunger (FEI Mark IV) at 100% humidity and plunge frozen in −172°C liquefied ethane. For the apo complex the grids are blotted for 2 s with a blot force of 10. Standard vitrobot filter paper ∅ 55/20mm, Grade 595 was used for blotting.

### Cryo-EM data collection and analysis

Micrographs of the Chd1-nucleosome apo complex were collected on an FEI-F20 electron microscope fitted with a field emission gun and a TVIPS F416 4k × 4k CMOS camera. The microscope was operated at 200 kV, spot size 1, with a C2 condenser aperture of 50 μm and an objective aperture of 100 μm diameter. Semi-automatic data acquisition was performed using EMtools (TVIPS). Micrographs of vitrified samples were recorded at a primary magnification of 68000 (pixel size 1.58Å /pixel) and with an approximate dose of 17-22 electrons/Å^2^. Data were processed and the cryoEM map of the final model was obtained with RELION 1.3 (Scheres, 2012). The volumes are visualised and models were fitted with Chimera (Pettersen et al., 2004).

For the Chd1 Δ 57-88 complex, grids were loaded into a Titan Krios electron microscope (FEI) for automated data collection over a period of 2 days using the EPU software (FEI). Images were recorded at a nominal magnification of ×59,000 on a Falcon II direct electron detector. In total 3200 micrographs were recorded using a −1.5/−3.5 μm defocus range with a total electron dose of 40 e^−^ perÅ^2^. Each micrograph contains 22 frames and a total exposure time of 0.9 sec with 2.1 e‐ per frame. All movies were corrected for beam-induced drift using the frame wise motion correction with Unblur (Brilot et al., 2012; Campbell et al., 2012). The contrast transfer function (CTF) parameters of each drift-corrected image were estimated using CTFFIND4 (Rohou and Grigorieff, 2015). Micrographs with large astigmatism, heavy contamination, or serious aggregation were discarded resulting in 2450 micrographs. RELION 1.4 (Scheres, 2012) was used for the rest of the data processing. About 5000 particles from 50 micrographs were first handpicked, then extracted and 2D classes were generated in Relion 1.4. These 2D classes were then used as a reference in Relion autopick routine and particles were picked from all 2450 micrographs. The autopicked particles were subsequently extracted and sorted using particle sorting routine. A first round of two-dimensional classification was performed to discard poorly averaging particles, resulting in a cleaned 197,602-particle data set. A three-dimensional classification was then performed using four classes and with low pass filtered nucleosome volume as the initial cryo-EM map; 148607 particles belonging to the best three-dimensional classes were selected, 3D auto refined and subjected to single particle movie correction (Rubinstein and Brubaker, 2015) and particle re-centering (Rawson et al., 2016). Subsequently a second round of three-dimensional classification was performed using 4 classes. This approach yielded an improved volume belong to single class calculated from 70,481 particles. This class was further subjected to an additional round of 3D classification using 3 classes and finer rotational angle. These classes were inspected in chimera and two similar classes were merged, resulting in 52208 particles. These were subsequently separated into 187 groups, on the basis of their refined intensity scale-factor, and subjected to a final three-dimensional refinement using the selected 60Å low-pass filtered three-dimensional class as a starting model. User defined soft edge solvent mask was provided for solvent inclusion during the refinement process. The density map was corrected for the modulation transfer function (MTF) of the Falcon detector. The final resolution after post-processing was 15Å, according to the 0.143 cut-off criterion. UCSF Chimera was used for automated rigid-body docking as well as to generate figures and videos(Pettersen et al., 2004). The cryo EM envelopes are deposited in the Electron Microscopy Data Bank (accession no. EMDB-3502 and xxxx) for release upon publication. Micrographs used to in this publication will be available upon publication via the image data repository http://idr-demo.openmicroscopy.org/about/ .

### Single Molecule FRET measurements

A biotin attached to the long linker of the nucleosome was used to bind nucleosomes to the surface of micro-fluidic chambers similiar to the attachment of transcription complexes (Andrecka et al., 2008). The chamber was inserted into a homebuilt TIRF microscope described earlier (Lewis et al., 2008) and smFRET data was recorded using alternating excitation (532 nm and 637 nm) at a frequency of 10 Hz for a total duration of 100-200 seconds (Treutlein et al., 2012). For all experiments, a smFRET buffer was used composed of 20 mM Tris/HCl pH 7.5, 0.5 mM EGTA, 50 mM NaCl, 3 mM MgCl_2_, 10% Glycerol, 2 mM DTT, 200 ng/μl BSA and freshly supplemented with the Oxygen Scavenger System (glucose oxidase, *Sigma-Aldrich*, 0.01 μg/μl final concentration; catalase, *Roche*; 1085 U/ml final concentration), D-(+)-glucose (*Sigma-Aldrich*, 4 % w/v) (Rasnik et al., 2006) and Trolox (*Fluka*, 1mM final concentration, illuminated with UV-light for 6 min prior to mixing with Oxygen Scavenger System) (Cordes et al., 2009). After acquiring 10-15 smFRET movies for mono-nucleosomes in the absence of Chd1, Chd1 was loaded to the nucleosomes at a concentration of 50 nM in smFRET buffer with or without 150 μM AMP-PNP (*Roche*) and approximately 15 more smFRET movies were recorded. Quantification of smFRET and HMM analysis was performed using a custom-written software (Sikor et al., 2013).

## Acknowledgements

We thank David Bhella and Saskia Hutten (University of Glasgow) and the cryo EM-facility in Edinburgh for assistance in screening cryo grids. Joanna Brown assisted with the preparation of cryo EM grids at the beginning of the project. High resolution data was collected at NeCEN with assistance from Ludo Renault and Christoph Diebolder. Sara Ten Have for assistance with analysis of crosslink MS data. This work was funded by Wellcome Senior Fellowship 095062, Wellcome Trust grants 094090, 099149 and 097945. ALH was funded by and EMBO long term fellowship ALTF 380-2015 cof-unded by the European Commission (LTFCOFUND2013, GA-2013-609409).

## Figure Legends

**Figure 1 figure supplement 1.**
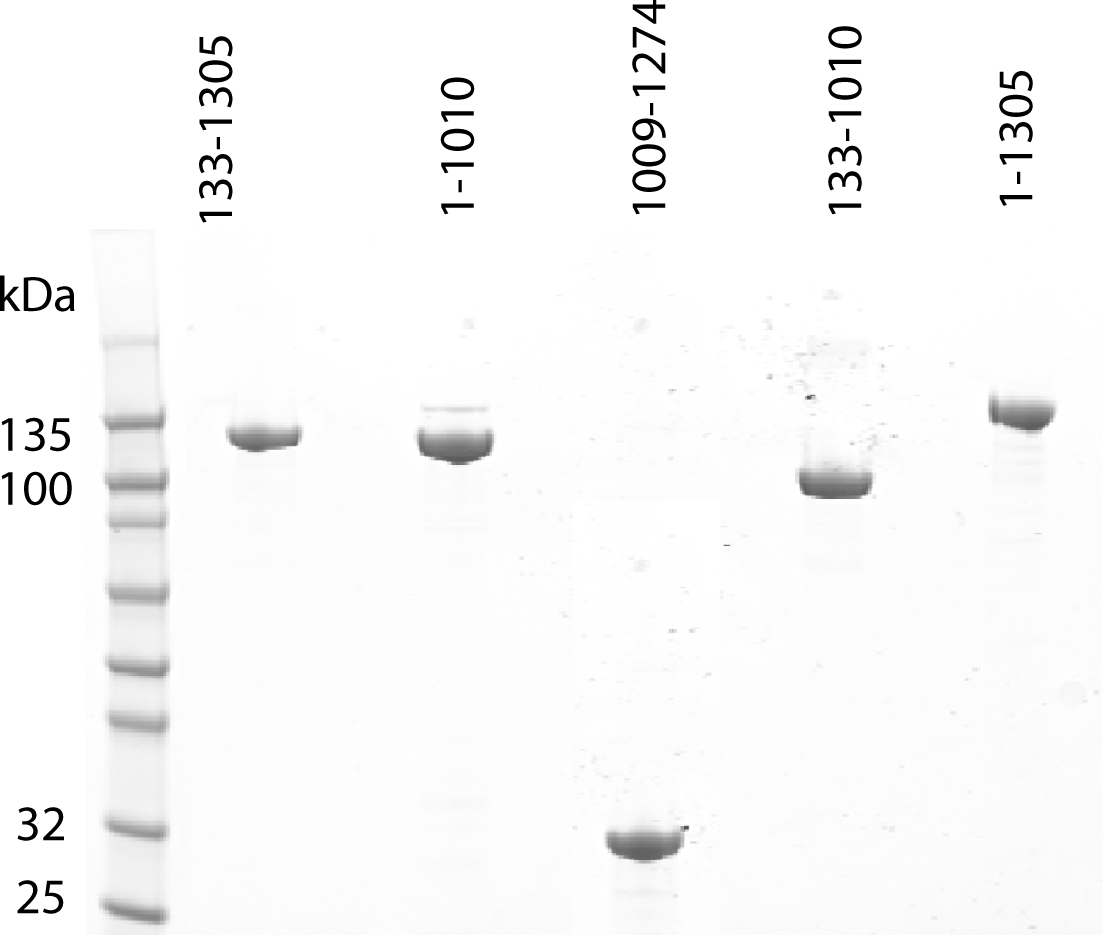
Purification of fragments of Chd1 protein. Fragments of purified Chd1 protein used in this study were loaded on an SDS PAGE gel and stained with coomassie. Numbers refer to the positions within the *S. cerevisiae* sequence.

**Figure 1 figure supplement 2.**
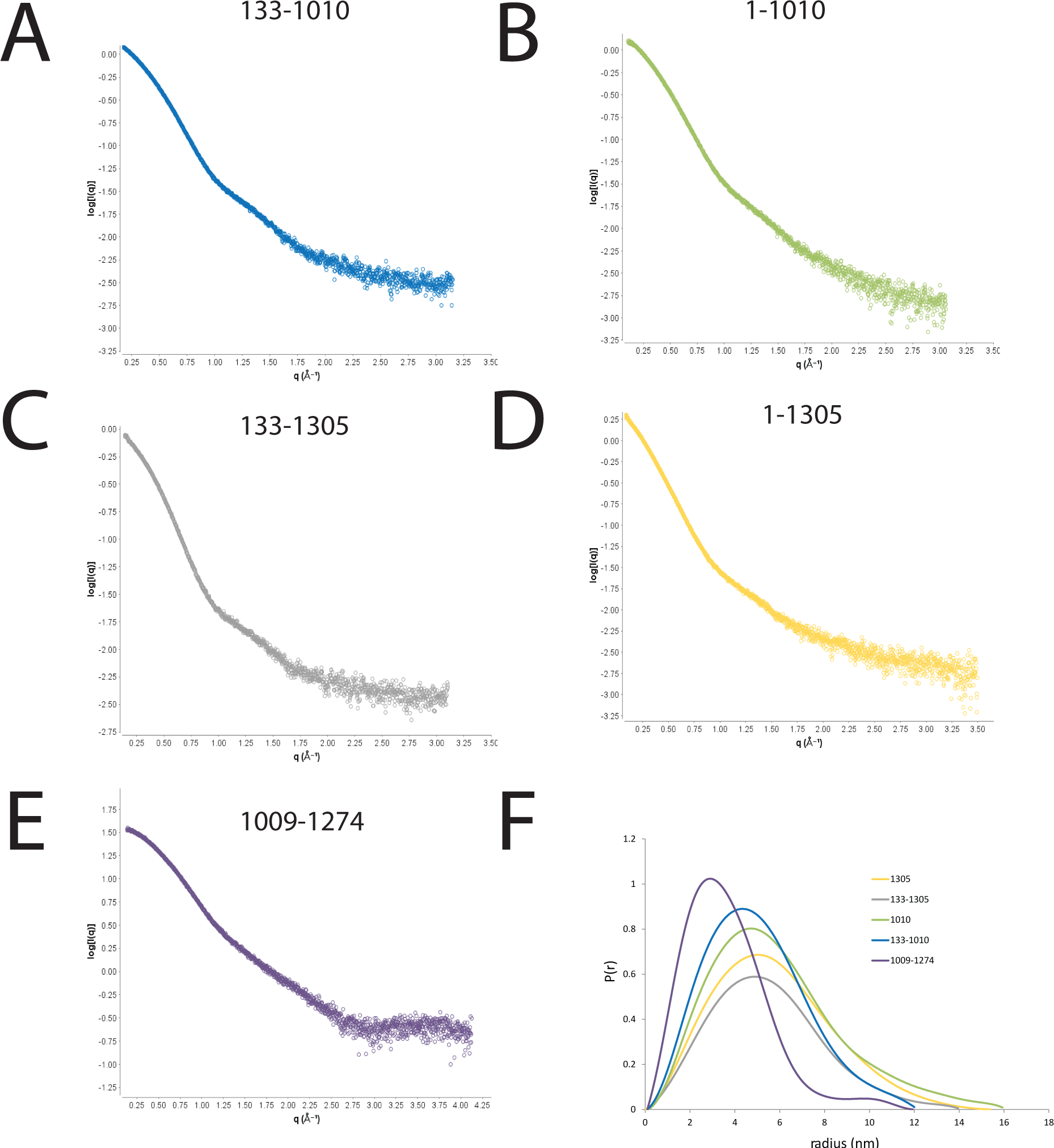
SAXS scattering curves for Chd1 fragments. A-E plots of the scattering intensity against the scattering vector for the following Chd1 fragments A 133-1010, B 1-1010, C 133-1305, D 1-1305, E 1009-1274. F shows the probability distribution for the radius of gyration for each fragment.

**Figure 2 figure supplement 1.**
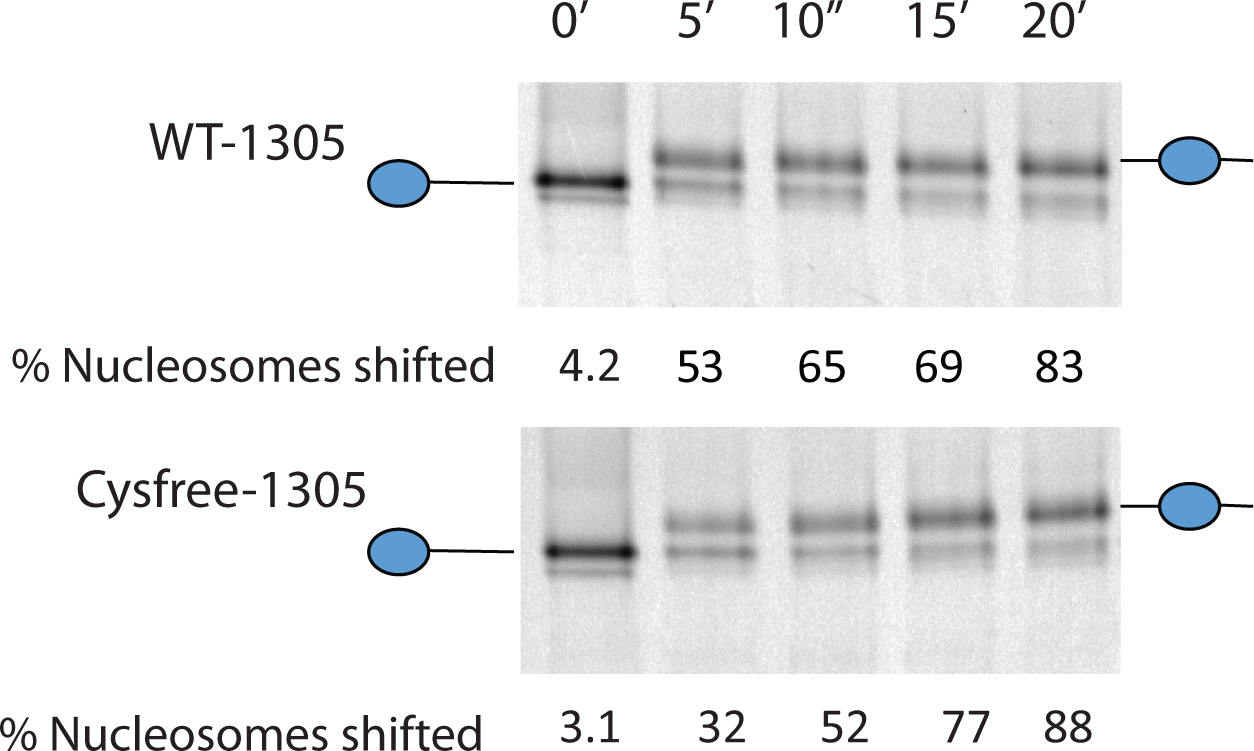
Activity of Chd1 following removal of native cysteines. Nucleosomes assembled onto the fragment 0W47 labelled with cy3 we used as a substrate for repositioning assays using Chd1 1-1305 (top panel) or Chd1 1-1305 with the six native cysteins mutated to serines. Both proteins cause a time dependent increase in the mobility of nucleosomes in the presence of ATP that is consistent with repositioning to a more central locations. Quantification of the proportion of nucleosomes repositioned indicates that the activities of the two proteins are comparable.

**Figure 2 figure supplement 2.**
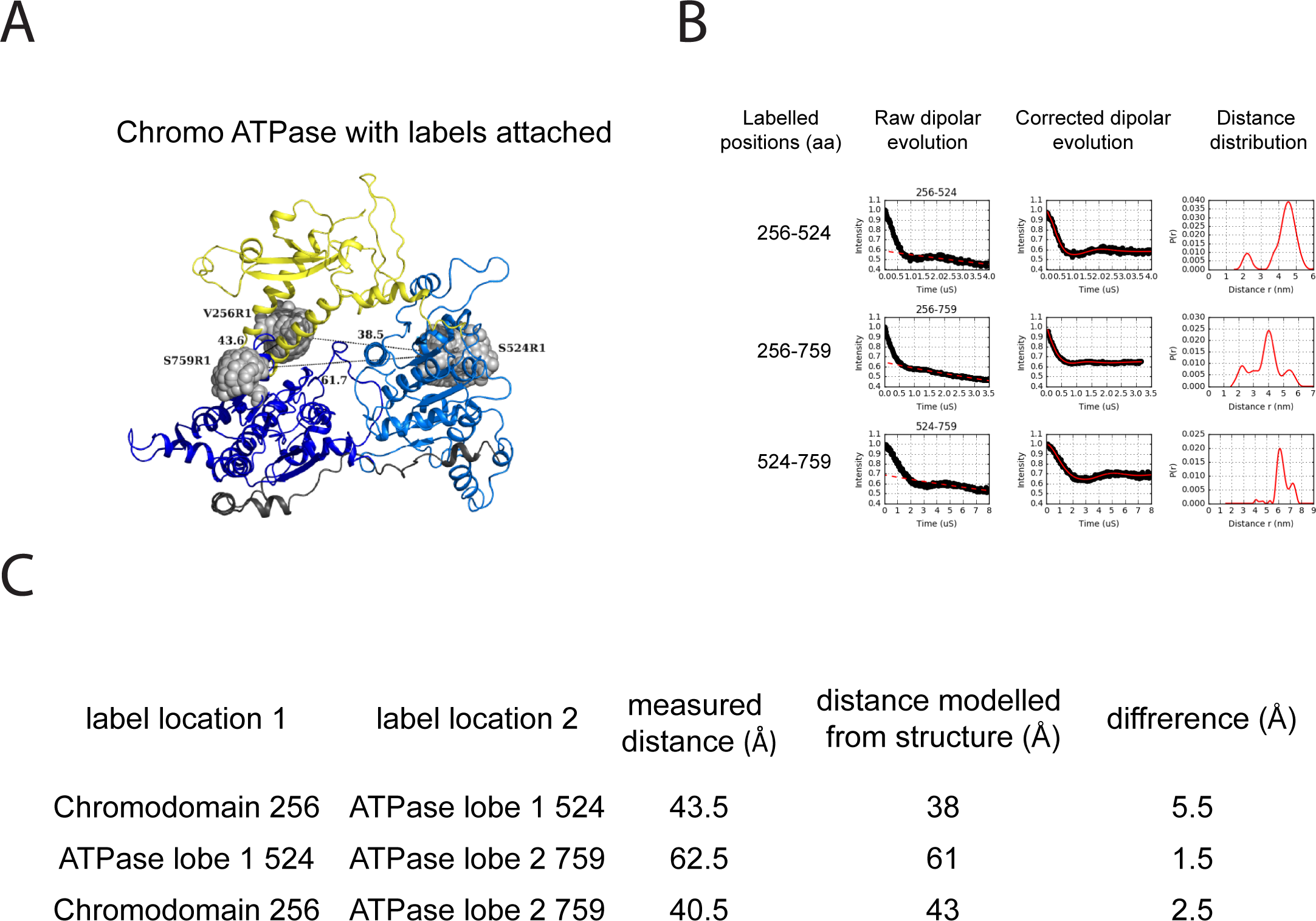
PELDOR measurements within the chromoATPase domains of Chd1. **A** Cartoon representation of the chromo helicase domain with molecular dynamics simulated conformers of MTSSL spin label nitroxide atom drawn as spheres. **B** PELDOR time traces performed between these labelling sites. **C** The measured distance determined experimentally is compared with the distance anticipated based upon modelling to the crystal structure of the chromoATPase domains and the difference between these measurements is shown.

**Figure 2 figure supplement 3.**
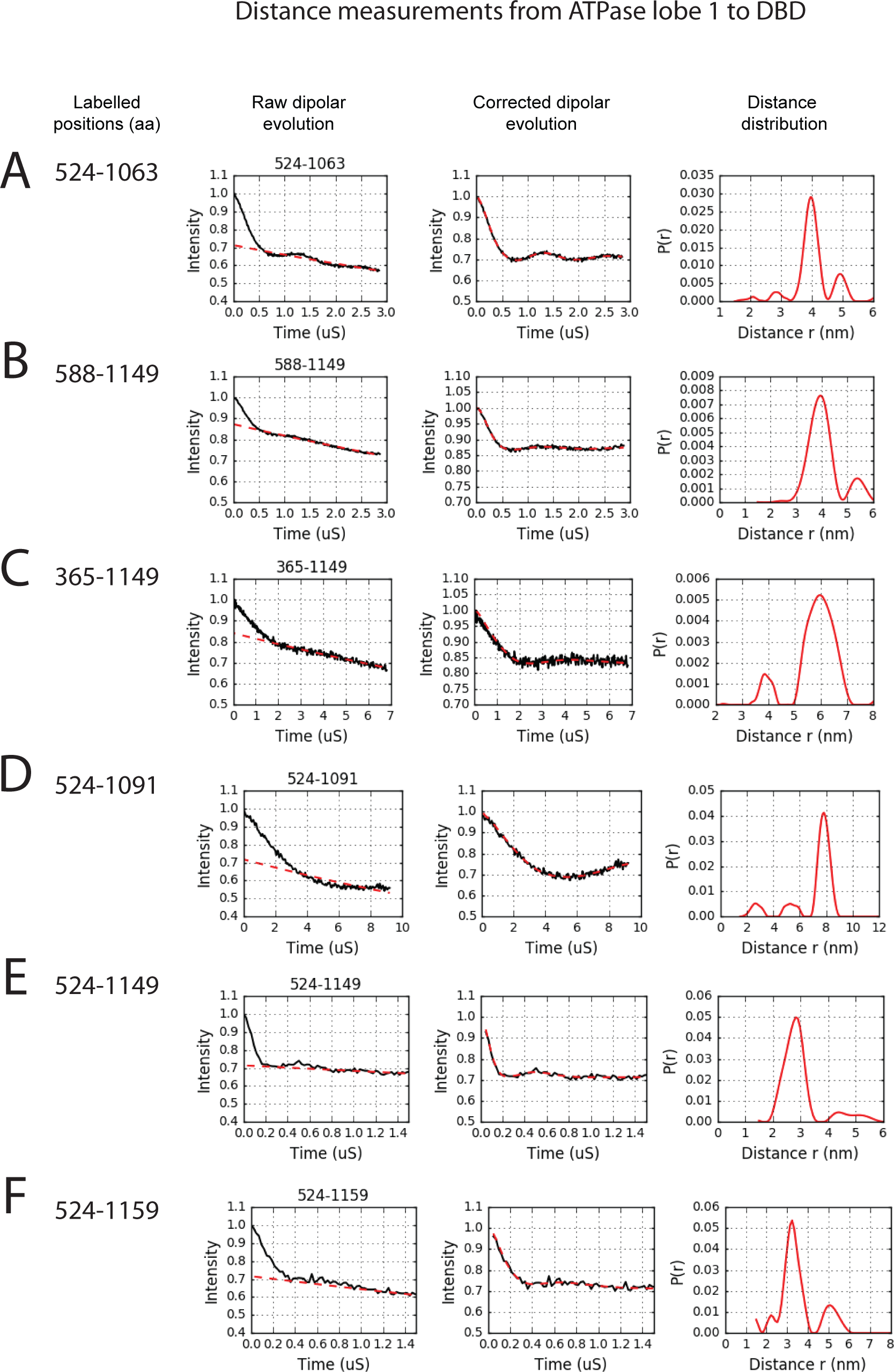
PELDOR measurements between ATPase lobe1 and the DNA binding domain. PELDOR data of the pair of spin labels attached to the indicated sites on the ATPase lobe 1 of the Chd1 (1-1305) molecule with indicated sites on the DNA binding domain. Raw dipolar evolution, background corrected evolution and distance distribution obtained after Tikhonov regularisation are shown for the indicated sites.

**Figure 2 figure supplement 4.**
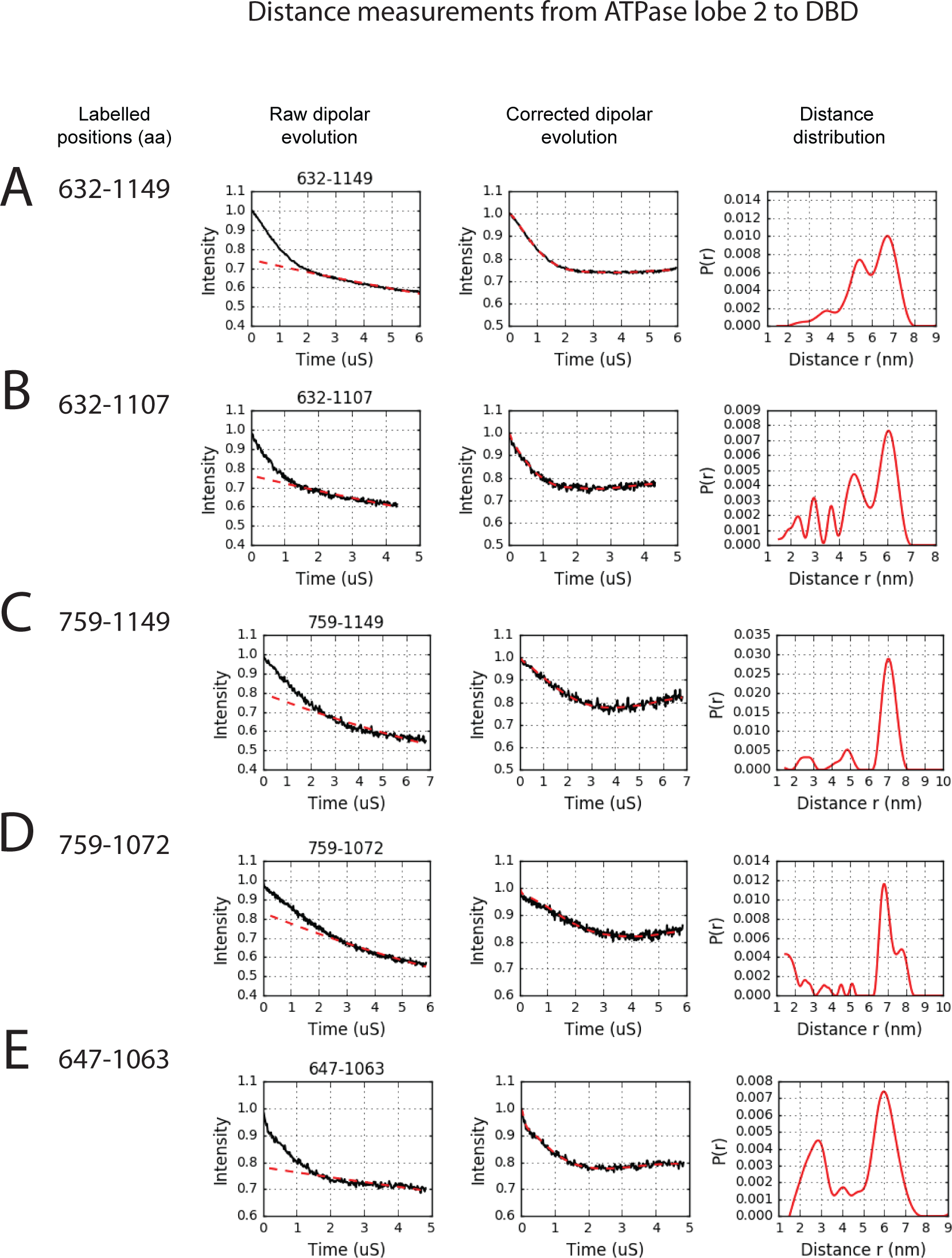
PELDOR measurements between ATPase lobe 2 and the DNA binding domain. PELDOR data of the pair of spin labels attached to the indicated sites on the ATPase lobe 2 of the Chd1 (1-1305) molecule with indicated sites on the DNA binding domain. Raw dipolar evolution, background corrected evolution and distance distribution obtained after Tikhonov regularisation are shown for the indicated sites.

**Figure 2 figure supplement 5.**
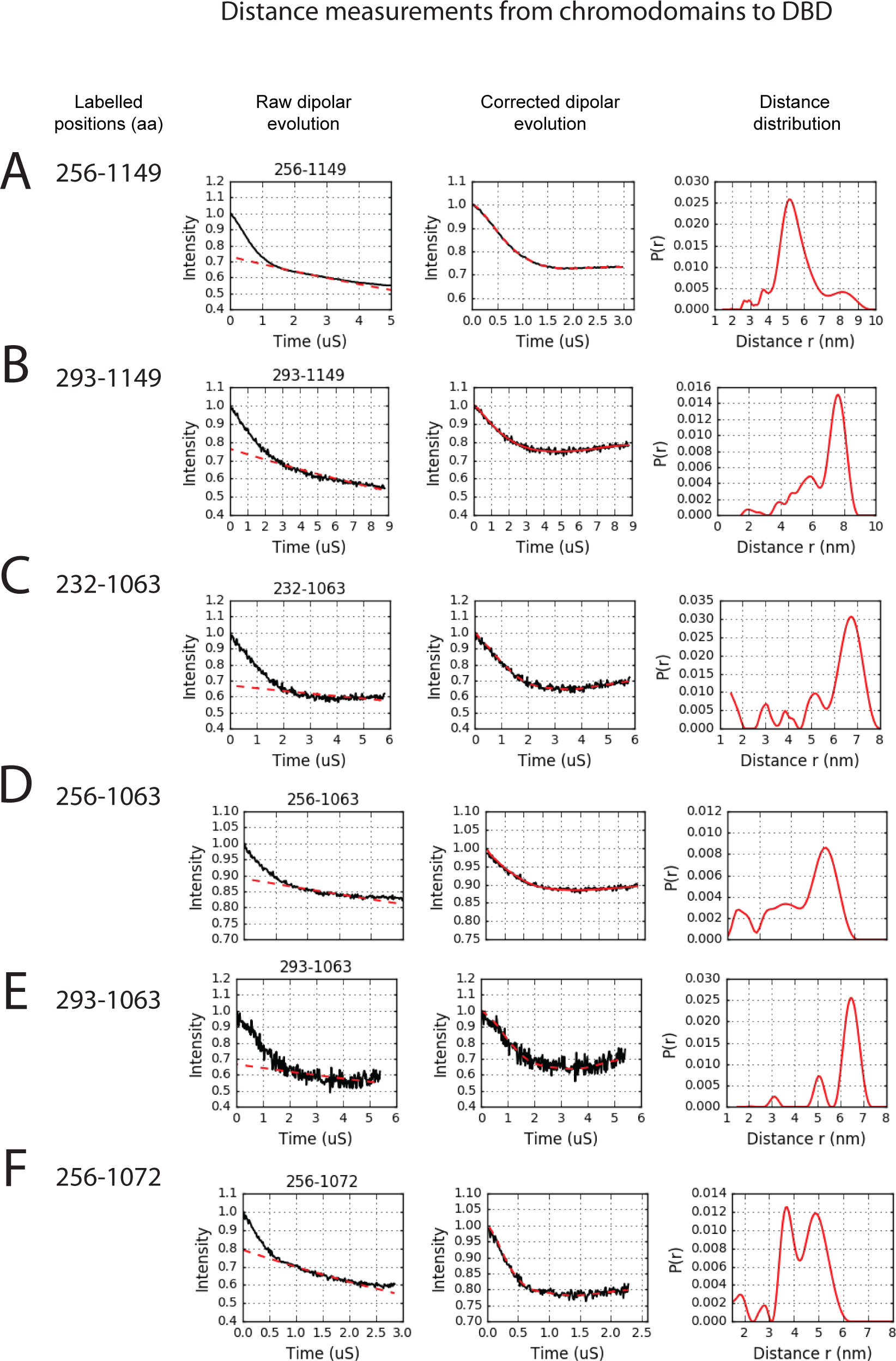
PELDOR measurements between chromodomains and the DNA binding domain. PELDOR data of the pair of spin labels attached to the indicated sites on the chromodomains of the Chd1 (1-1305) molecule with indicated sites on the DNA binding domain. Raw dipolar evolution, background corrected evolution and distance distribution obtained after Tikhonov regularisation are shown for the indicated sites.

**Figure 2 figure supplement 6.**
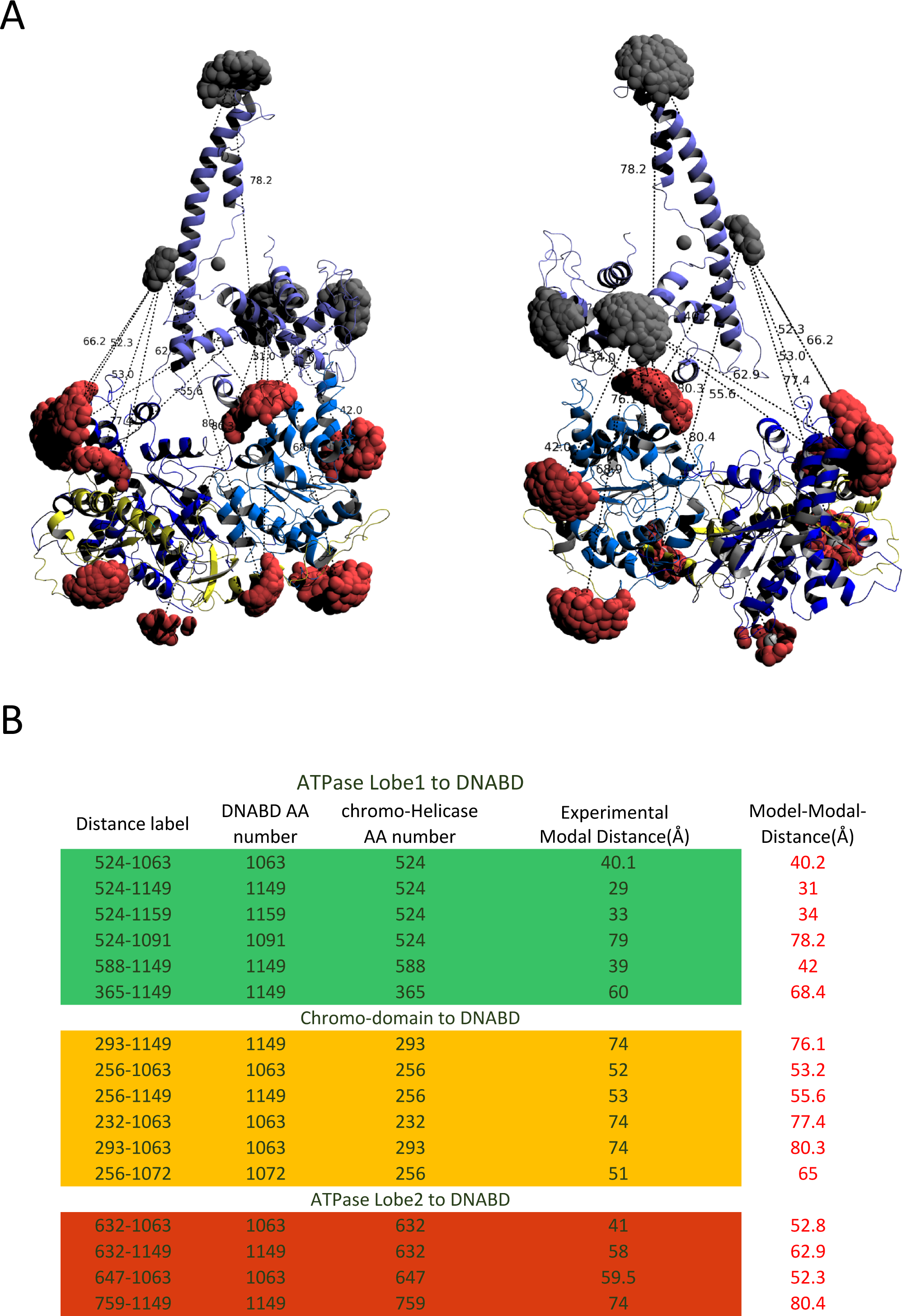
Modelled orientation of chromoATPase and DNA binding domain. **A** Cartoon representation of the chromo helicase and DNA binding domain with measured distances indicated. **B** The measured distances determined experimentally are compared with the distances anticipated based upon the averaged model obtained in Figure 2E.

**Figure 2 figure supplement 7.**
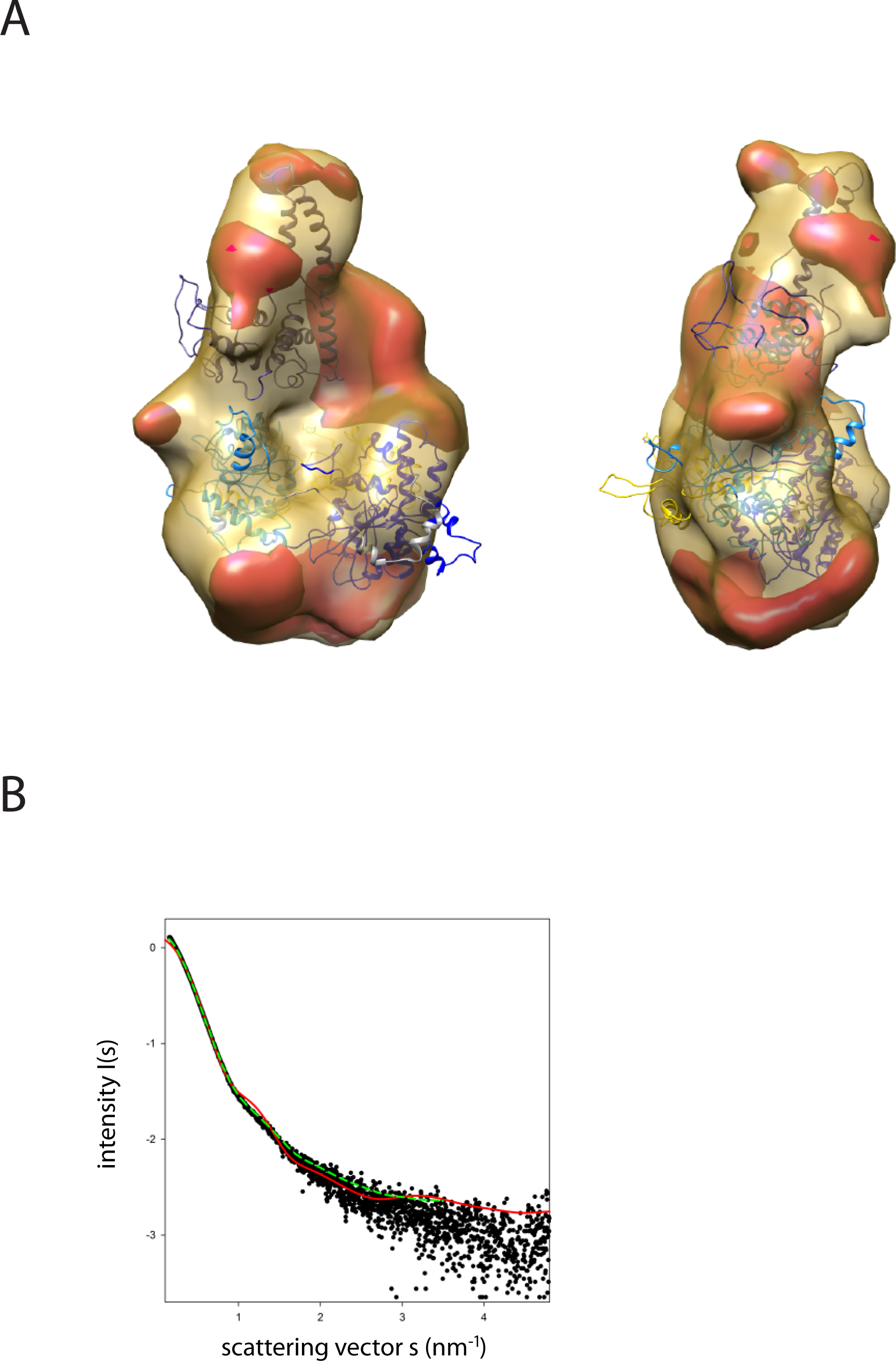
Fit of PEDOR model for Chd1 into SAXS volume. **A,** Ab-initio generated SAX volume correspond to the 1305 protein is shown in two orientations. The structure fitted into volume is the model generated from rigid body docking of known crystal structures using sparse PELDOR distance distribution measurements. The difference volume coloured in red-orange indicates the unresolved structures of Chd1 protein (1-175aa in the N-terminal and 932-1010). **B**, Comparison of the theoretical scattering curve generated by the alignment of the DNA binding domain and chromoATPase using the PEDLDOR model (solid red) X2=11.5 and following inclusion of N-terminal and linker regions as dummy residues (dashed green) X2=2.55 with the experimentally obtained scattering from Chd1 1-1305 (black spots).

**Figure 5 figure Supplement 1.**
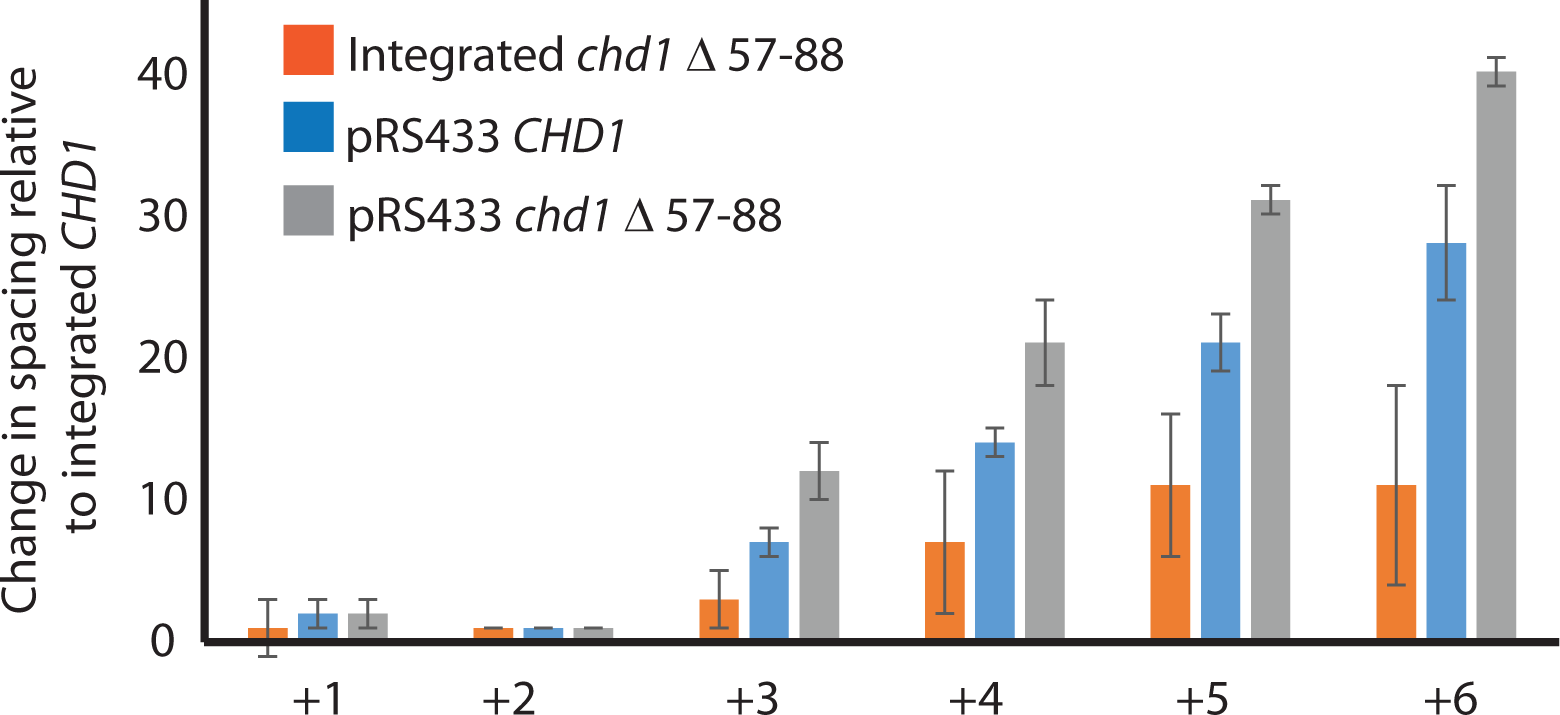
Changes in nucleosome positioning following reintroduction of different Chd1 constructs. Quantitative changes in nucleosome spacing in coding in strains expressing CHD1 (wt/(Δ57-88) at low (int-integrated) or high (Ectopically from multicopy plasmid pRS423) levels. The shift in nucleosome positions (+1 to +6) in coding region was calculated relative to integrated integrated-*CHD1* and plotted. The progressive increase in the positioning defect for more distal nucleosomes indicates a difference in average nucleosome spacing. Error bars indicate the standard deviation from three independent repeats.

**Figure 5 figure Supplement 2.**
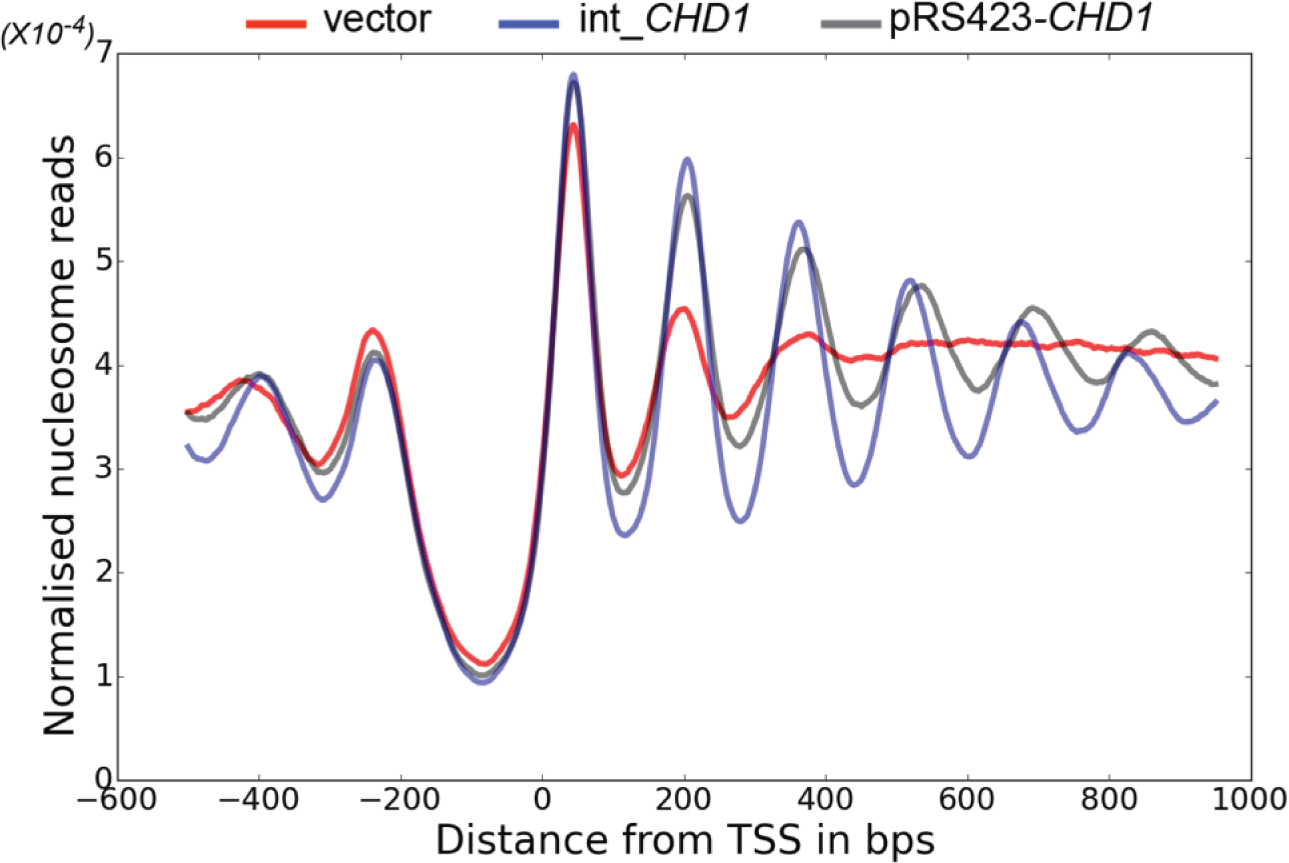
Changes in nucleosome positioning following reintroduction of *CHD1* at low or high copy number. Changes in nucleosome spacing in coding regions were measured in *Isw1Δchd1Δ* strains transformed with a control vector (vector, red) or in which *CHD1* was reintroduced integrated onto the chromosome in single copy (int-*CHD1, blue*) or at high copy on the multi-copy plasmid pRS423 (pRS423-*CHD1, green*). TSS-aligned nucleosome occupancy profiles were plotted for the above strains show increased spacing in the strains with higher levels of *CHD1*.

**Figure 5 figure Supplement 3.**
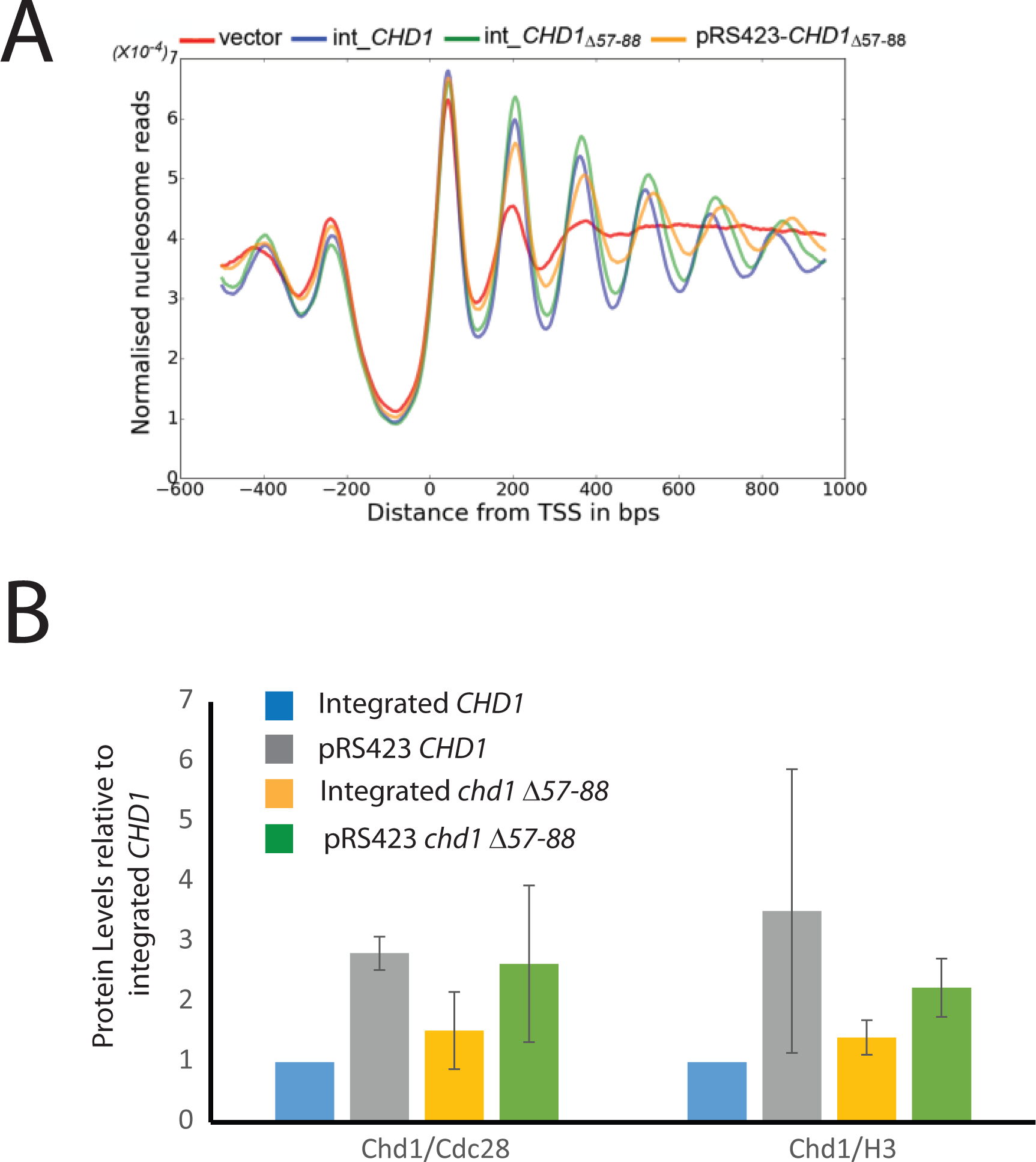
Changes in nucleosome positioning following reintroduction of chd1 *CHD1Δ57-88* at low or high copy number. AChanges in nucleosome spacing in coding regions were measured in strains expressing *CHD1Δ57-88* re-integrated in single copy at the *CHD1* locus (int_CHD1Δ57-88, green) or at higher copy number on pRS423 (pRS423_CHD1Δ57-88, orange). Vector only (red) and *CHD1* wt reintroduced integrated onto the chromosome in single copy (int-*CHD1, blue*) from Figure 5 are also shown for comparison. TSS-aligned nucleosome occupancy profiles are plotted for the above strains showing increased spacing in strain with *CHD1Δ57-88* compared to a strain in which wild type CHD1 has been integrated at single copy (int_ *CHD1).* Replacement on a high copy plasmid results in 2 to 4-fold increased Chd1 expression. BQuantitative measurement of protein levels in strains expressing *CHD1wt* or *CHD1Δ57-88* at low (int) or high (pRS423) copy number. For all strains described in A, Chd1 was N-terminally flag tagged and anti-FLAG western blotting used to plot protein levels relative to int-*CHD1*wt. Quantitation is plotted following normalisation against total histone H3 and Cdc28 as indicated. Error bars indicate standard error from 3-4 measurements.

**Figure 7 figure supplement 1.**
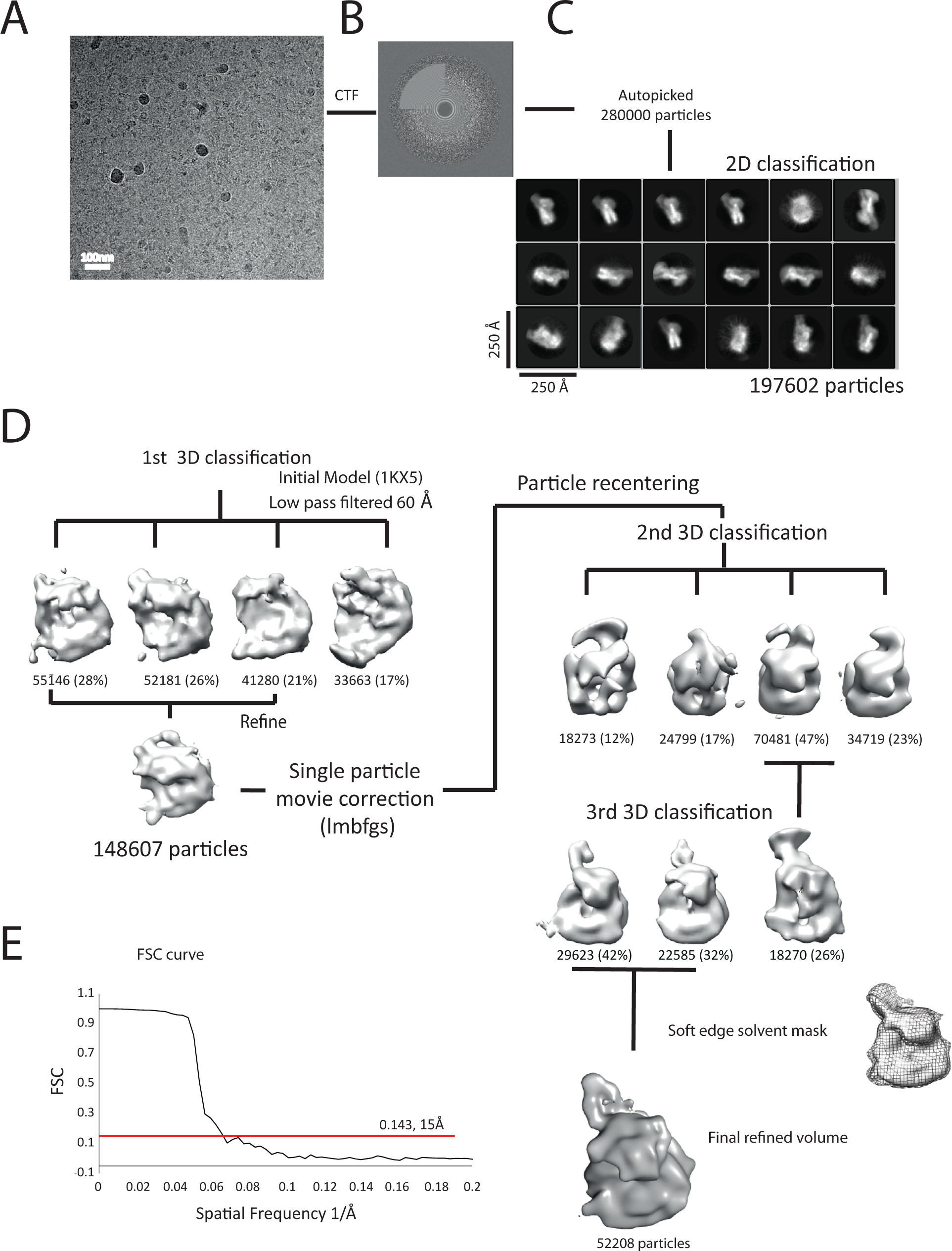
Overview of the cryoEM-data. **A**, Representative micrograph of frozen hydrated nucleosome-Chd1 complex. **B**, The CTFFIND4 output generated from the micrograph shown in (A). The experimentally observed Thon rings align with the predicted Thon rings (shown as a quadrant). **C**, Two-dimensional class averages of the CTF corrected auto picked particles is shown. Many of the classes have a visible attachment adjacent to the nucleosome. **D**, Workflow of the three-dimensional classification and structure refinement. In the first 3D classification, the crystal structure of the nucleosome (1KX5) was converted to a low pass filtered volume map and used as reference map. The particle numbers in each class are indicated. For the single particle movie correction lmbfgs (Limited Memory Broyden–Fletcher–Goldfarb–Shanno) algorithm was used. The two major classes from the third 3D classification were subject to refinement following application of a solvent mask applied using Relion 1.4. The final refined volume has 52208 particles and estimated resolution of 15Å. **E**, Gold standard Fourier-Shell correlation and resolution using the 0.143 criterion.

**Figure 7 supplement 2.**
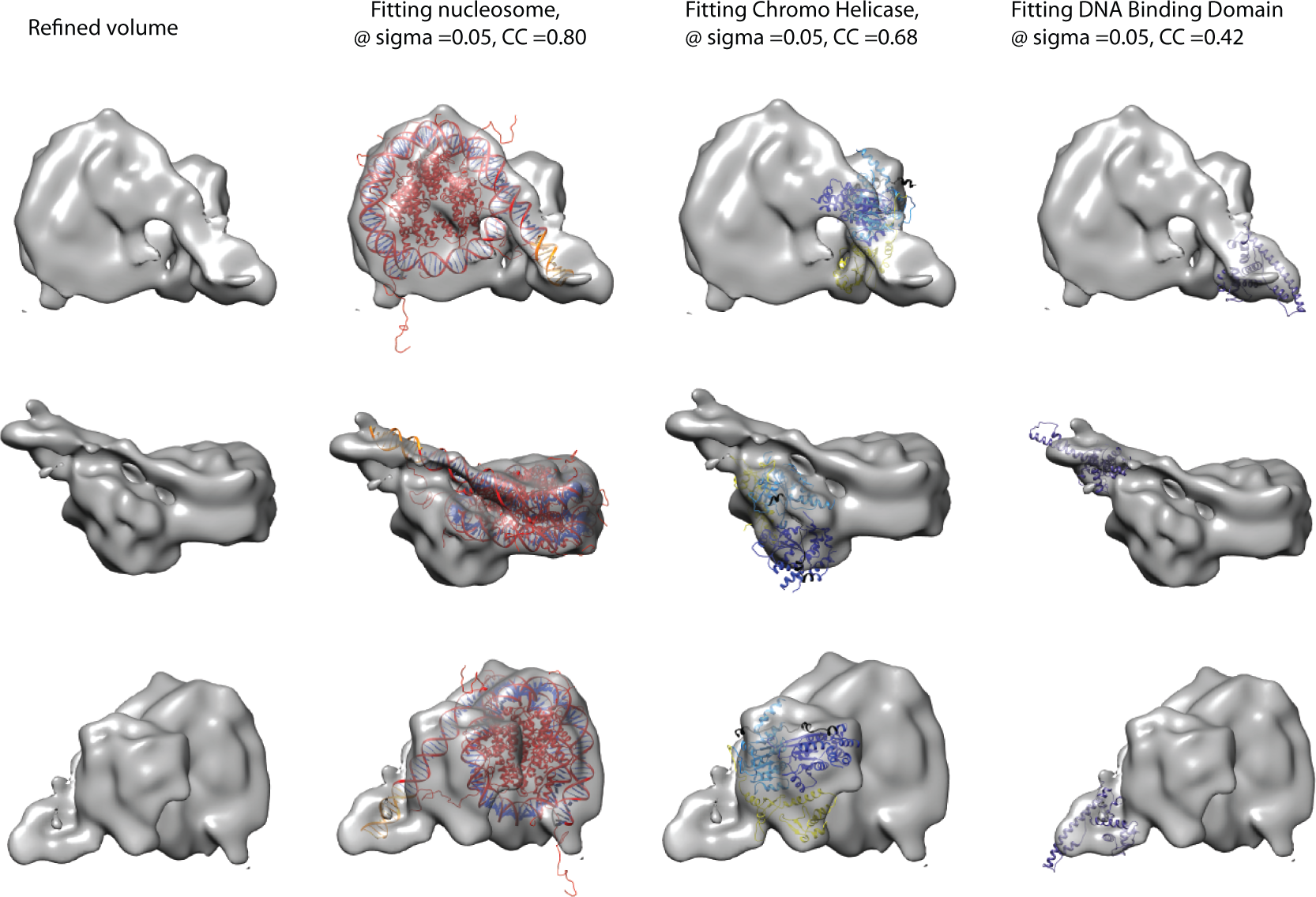
Fitting of nucleosome and Chd1 crystal structures into cryoEM map. Three views of the final refined electron density map of the nucleosome-Chd1 complex and the docked crystal structures of nucleosome (1KX5) with the extended linker DNA, chromo-helicase (3MWY) and the DNA binding domain are shown. The correlation coefficient calculated for the rigid body fit @ 0.05 sigma level is indicated.

**Figure 7 supplement 3.**
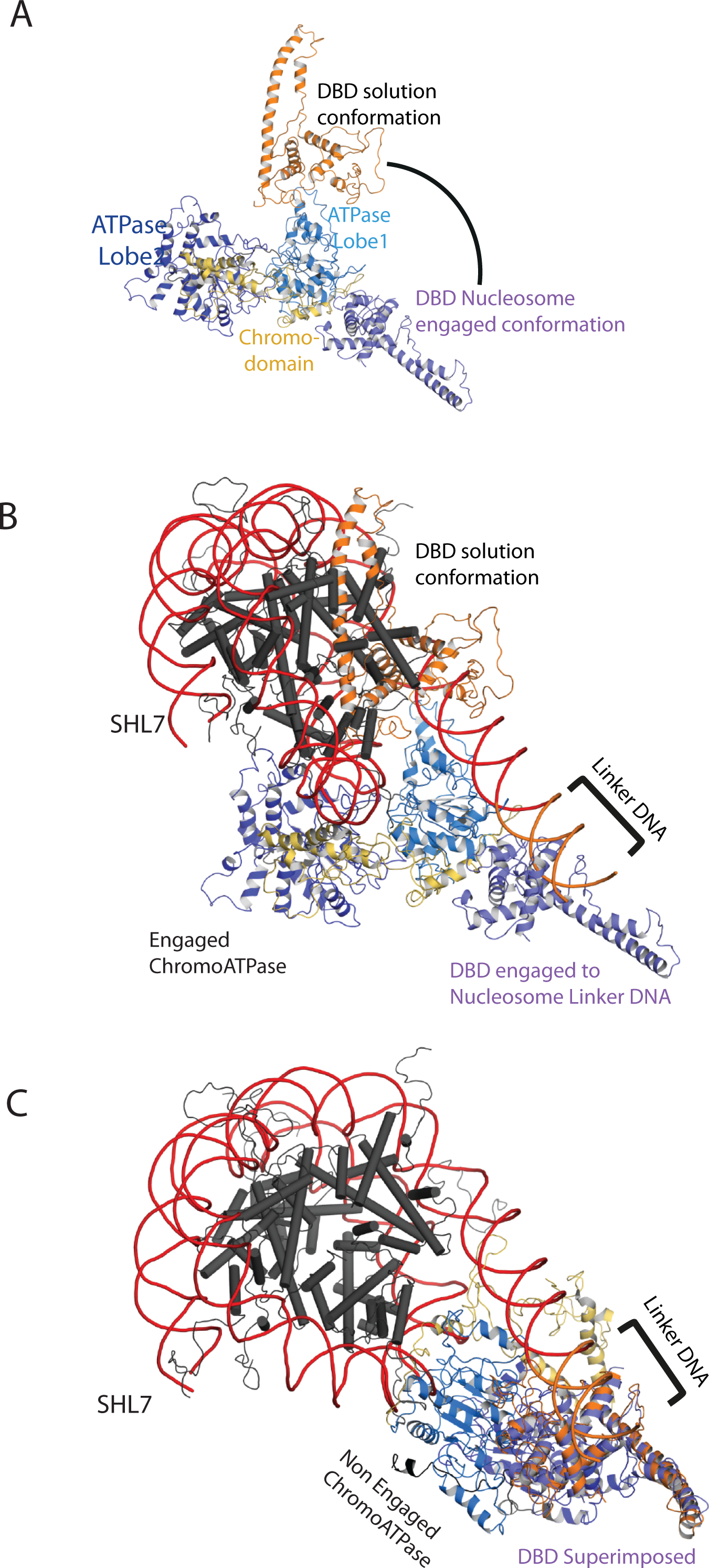
Comparison of DNA binding domain orientation in solution structure and when engaged with nucleosomes. (A) Solution of the structural model for Chd1 in solution derived from Figure 2 is superimposed on to Chd1 in the conformation observed when it is engaged with nucleosomes. The DBD domain from the solution model is coloured in orange and the nucleosome bound state in deep blue. (B) The solution model for Chd1 is docked onto a nucleosome using the location of the ATPase domains within the engaged state as a reference point. This conformation is not feasible as there are major clashes with the histone octamer. (C) The solution model for Chd1 is docked onto a nucleosome using the location of DBD in the nucleosome engaged state as a reference point. This conformation is related to that observed for Chd1 in the apo state (Figure 6).

**Figure 7 supplement 4.**
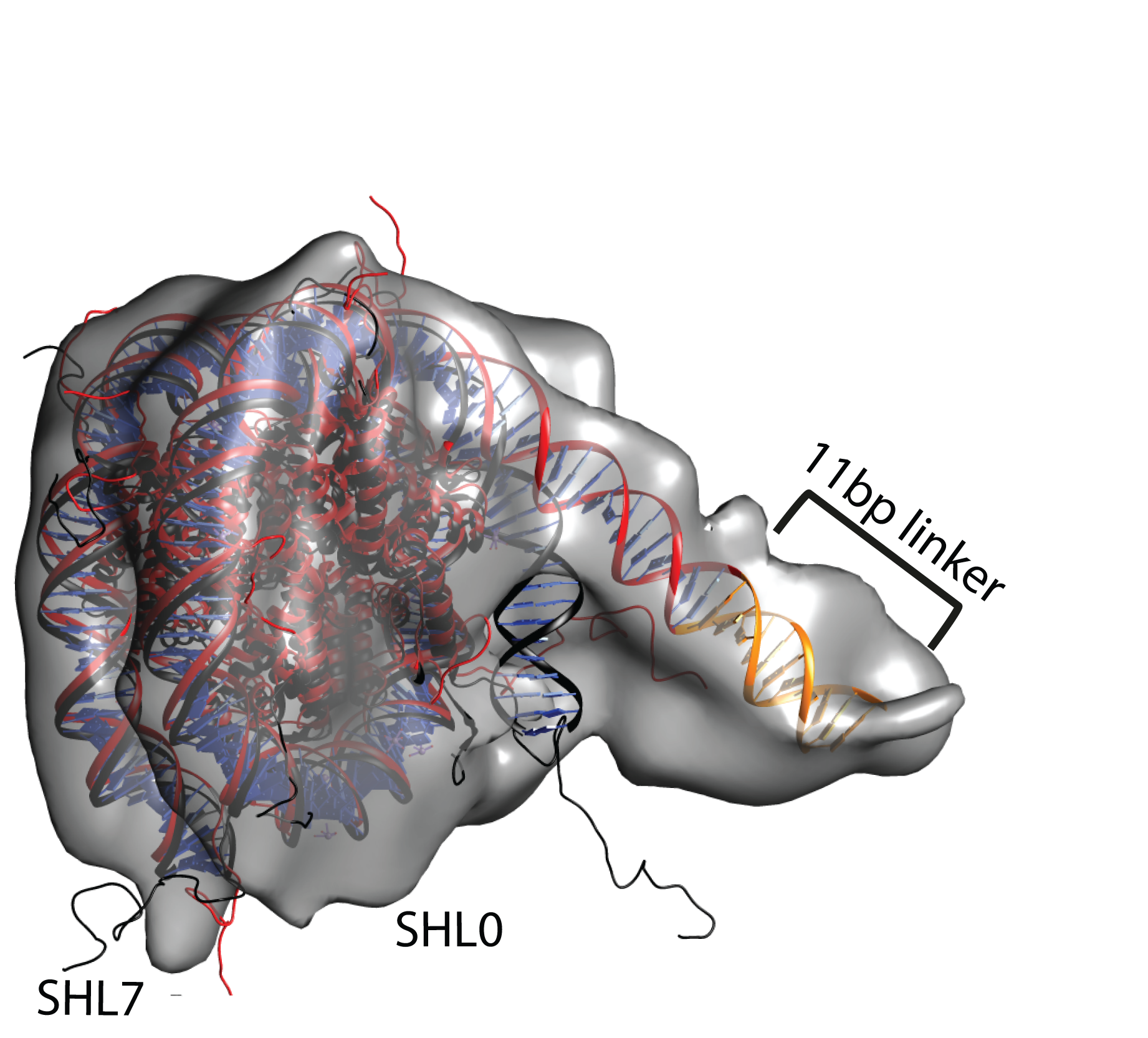
Unravelling of nucleosomal DNA adjacent to the bound linker. Refined cryoEM map of nucleosome-Chd1 complex is shown in semi-transparent with docked nucleosome (1KX5) crystal structure. The extended 11-bp linker DNA on one side of the nucleosome as defined in the electron density map is coloured in orange. The modelled nucleosome in the cryoEM map (red) is compared with the nucleosome crystal structure (Black). The outer turn of a fully wrapped nucleosome protrudes out of the envelope supporting unwrapping to accommodate the altered trajectory. The nucleosome dyad axis is marked as SHL0 and the linker free side of the nucleosome is marked as SHL7.

**Figure 8 figure supplement 1.**
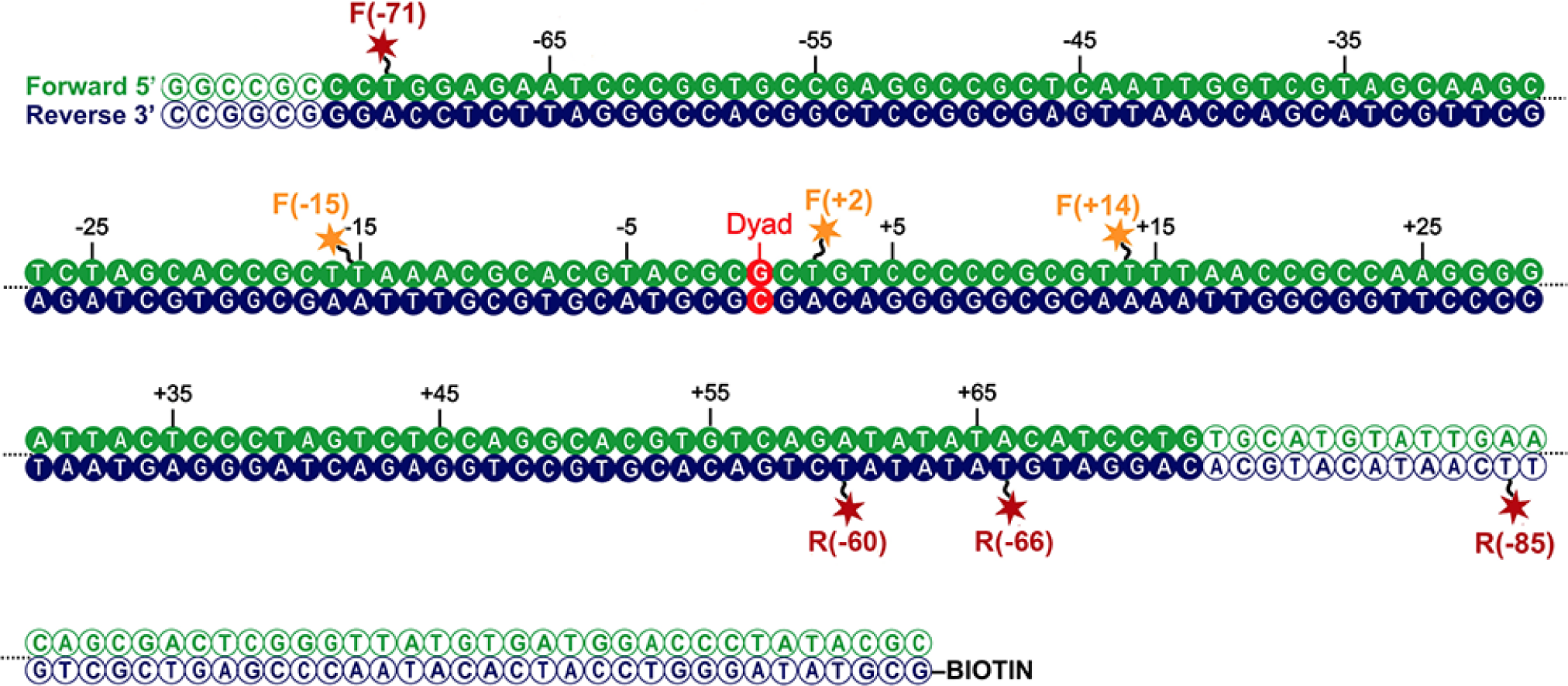
Map indicating locations to which fluorescent dyes are attached. Schematic illustration of the nucleosomal DNA and designed label positions, depicted as stars (red, acceptor dye Alexa647; yellow, donor dye Tamra). Nucleotides marked as open circles constitute the linker DNA, whereas nucleotides shown as filled circles are part of the 601 sequence and therefore constitute DNA inside the nucleosome core.

**Figure 8 figure supplement 2.**
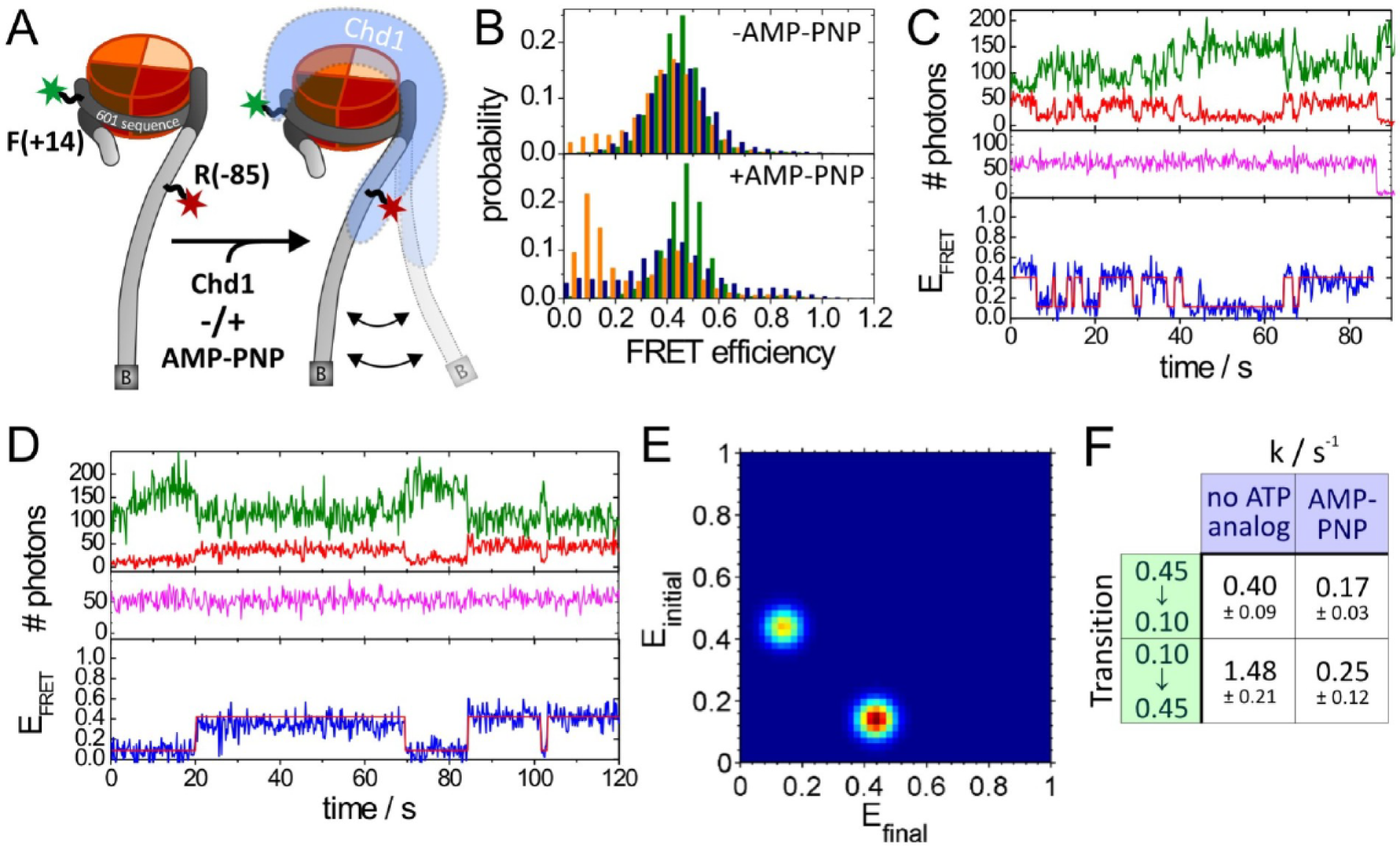
Quantitative smFRET analysis of DNA linker dynamics introduced by Chd1 binding. (**A**) Cartoon illustrating the smFRET measurement of nucleosomes labelled at position R-85 with Alexa647 and at position F+14 with Tamra. Observed dynamics of the linker DNA (see panel **C-F**) in the presence of Chd1 (shaded blue shape) are indicated. (**B**) FRET efficiency histograms of nucleosomes alone (green), nucleosomes in the presence of 50nM Chd1 showing static FRET (blue) as well as dynamic FRET trajectories (orange). The upper panel shows data of the measurements without ATP analog and the lower panel shows data of the measurements with 150μM AMP-PNP. (**C**) Exemplary dynamic FRET time trace in the presence of Chd1 without AMP-PNP. Time trace of donor fluorescence (green, upper panel) and acceptor fluorescence upon green excitation (red, upper panel) are shown together with the acceptor fluorescence upon direct excitation at 637 nm (magenta, middle panel). In the lower panel, the computed FRET efficiency (blue) is presented together with the time dependent transitioning between different FRET states as identified by HMM analysis (red). (**D**) Exemplary dynamic FRET time trace in the presence of Chd1 and AMP-PNP. Same color coding as in panel **C**. (**E**) Exemplary transition density plot (TDP) for transitions resulting from HMM analysis of a total of 184 dynamic traces of nucleosomes in the presence of Chd1 alone showing 791 transitions. (**F**) Table presenting transition rates and their standard deviations in extracted from a mono-exponential fit to the cumulative distribution of dwell times of each transition.

